# Molecular determinants of Karyopherin-β2 chaperone and disaggregation activity

**DOI:** 10.1101/2022.09.14.508025

**Authors:** Charlotte M. Fare, Kevin Rhine, Andrew Lam, Sua Myong, James Shorter

## Abstract

Karyopherin-β2 (Kapβ2) is a nuclear-import receptor that recognizes proline-tyrosine nuclear localization signals (PY-NLSs) of diverse cytoplasmic cargo for transport to the nucleus. Kapβ2 cargo include several disease-linked RNA-binding proteins (RBPs) with prion-like domains (PrLDs), such as FUS, TAF15, EWSR1, hnRNPA1, and hnRNPA2. These RBPs with PrLDs are linked via pathology and genetics to debilitating degenerative disorders, including amyotrophic lateral sclerosis (ALS), frontotemporal dementia (FTD), and multisystem proteinopathy (MSP). Remarkably, Kapβ2 prevents and reverses aberrant phase transitions of these cargo, which is cytoprotective. However, the molecular determinants of Kapβ2 that enable these activities remain poorly understood, particularly from the standpoint of nuclear-import receptor architecture. Kapβ2 is a superhelical protein comprised of 20 HEAT repeats. Here, we design truncated variants of Kapβ2 and assess their ability to antagonize FUS aggregation and toxicity in yeast and FUS condensation at the pure protein level and in human cells. We find that HEAT repeats 8-20 of Kapβ2 recapitulate all salient features of Kapβ2 activity. By contrast, Kapβ2 truncations lacking even a single cargo-binding HEAT repeat display reduced activity. Thus, we define a minimal Kapβ2 construct for delivery in adeno-associated viruses as a potential therapeutic for ALS/FTD, MSP, and related disorders.

## Introduction

Karyopherin-β2 (Kapβ2; also known as Transportin 1) is a nuclear-import receptor (NIR) that transports proteins bearing a proline-tyrosine nuclear localization signal (PY-NLS) from the cytoplasm to the nucleus (1–4). Kapβ2, like other NIRs, promotes nuclear import based on the presence of an asymmetric Ran gradient (3,5). Ran is a small GTPase, and its GTPase-activating protein (GAP), RanGAP1, is restricted to the cytoplasm, whereas its guanine-nucleotide-exchange factor (GEF), RCC1, is localized to the nucleus (3, 5). Thus, Ran-GDP is predominant in the cytoplasm, whereas Ran-GTP is predominant in the nucleus (3,5,6). Kapβ2 has a low affinity for Ran-GDP, which permits Kapβ2 interactions with cargo in the cytoplasm (3, 7). Once in the nucleus, Kapβ2 is engaged tightly by Ran-GTP, which elicits a conformational change in Kapβ2 that releases cargo and liberates Kapβ2 for repeated rounds of nuclear import (3,7–10).

The best-characterized class of cargo recognized by Kapβ2 are proteins bearing a PY-NLS (1,2,11–14). The PY-NLS is defined by three distinct epitopes that each contribute to cargo-recognition. A hydrophobic or basic motif is epitope 1, which is followed by a R-X_2-5_-PY motif where R-X_2-5_ is epitope 2 and the C-terminal PY is epitope 3 (1,12,13). Kapβ2 was initially described with respect to its role in nuclear import (7,10,11,15). More recently, Kapβ2 has been defined as a potent molecular chaperone that can prevent and reverse aberrant phase transitions of its specific cargo (16–24).

The newly appreciated chaperone and disaggregation activity of Kapβ2 could have therapeutic utility (6,25–28). Indeed, several of the cargo recognized by Kapβ2 are prominent RNA-binding proteins (RBPs) with prion-like domains (PrLDs), which are connected to neurodegenerative disease via both pathology and genetics, including FUS, hnRNPA1, hnRNPA2, TAF15, and EWSR1 (8,10,11,15,16,29–34). In particular, mutations in genes encoding RBPs with PrLDs can cause amyotrophic lateral sclerosis (ALS), frontotemporal dementia (FTD), multisystem proteinopathy (MSP), oculopharyngeal muscular dystrophy, hereditary motor neuropathy, and related disorders (35–41). In postmortem brain tissue of ALS/FTD patients, nuclear RBPs with PrLDs have been found to be mislocalized and aggregated in the cytoplasm of degenerating neurons (37,38,42–48). The manner by which cytoplasmic mislocalization of specific RBPs with PrLDs elicits neurodegeneration has two dimensions: (1) RBPs play an essential role in RNA metabolism (28,44,49–51), and their cytoplasmic mislocalization may elicit loss-of-function phenotypes that contribute to toxicity; and (2) the insoluble cytoplasmic forms of these RBPs may be intrinsically toxic (42,52–57), indicating a gain-of-function mechanism of toxicity.

Despite the potential risk of forming toxic assemblies, the ability for RBPs to self-assemble into phase-separated states also has biological utility (28,58–60). RBPs with PrLDs can mediate multivalent intermolecular contacts, allowing them to self-associate and condense into phase-separated bodies (28,55,58,61–64). In particular, RBPs with PrLDs like FUS and hnRNPA1 partition into stress granules (SGs), phase-separated membraneless organelles that form in the cytoplasm in response to stress (16,28,39,55,62,65–68). Dysregulation of SGs has been implicated in the appearance of cytoplasmic aggregates, which are associated with the development of ALS/FTD and MSP (31,39,55,62,69–71). Indeed, liquid condensates formed by RBPs with PrLDs can mature into pathological solid phases comprised of cross-β fibrils that are associated with disease, and this process is often accelerated by disease-linked mutations (20,55,69,72). Ensuring that RBPs do not get trapped in pathological phases is a problem that all cells must solve (25, 26).

Cells have evolved agents to dissolve aberrant protein states, including NIRs (25, 26). *In vitro*, Kapβ2 prevents and reverses the formation liquid-like condensates, dense hydrogels, and solid-like fibrils by RBP cargo, including FUS and hnRNPA1 (16-22,24). Moreover, Kapβ2 performs these activities *in vivo* to antagonize the formation of cytoplasmic RBP aggregates, restore functional RBPs to the nucleus, and mitigate disease-linked RBP toxicity (6,16,27). Thus, by combining chaperone and nuclear-import activities, Kapβ2 antagonizes cytoplasmic RBP aggregation and the loss of RBP function in the nucleus (16).

In addition to recognizing cargo with a PY-NLS, Kapβ2 exhibits NLS-independent chaperone activity (27, 73). For example, Kapβ2 can antagonize aberrant phase transitions and interactions of ALS/FTD-associated non-native poly(GR) and poly(PR) dipeptide-repeat proteins (DPRs) produced by repeat-associated non-AUG (RAN) translation of the pathogenic G_4_C_2_ hexanucleotide repeat expansion in the *C9ORF72* gene (27,73–76). Kapβ2 mediates this chaperone activity by interacting with arginine residues, a strategy that is employed by a host of NIRs to chaperone a network of native and mutant clients (22, 23).

Kapβ2 chaperone activity is limited by its affinity for cargo. Kapβ2 typically recognizes cargo with nanomolar affinity (1,12,14,23). Two tryptophan residues, W460 and W730, on the cargo-binding surface of Kapβ2 play a critical role in cargo recognition, as they participate in hydrophobic sidechain stacking interactions with the PY-NLS (1). Importantly, a mutant form of Kapβ2, Kapβ2^W460A:W730A^, which has reduced affinity for PY-NLS-bearing cargo displays reduced chaperone activity (1, 16).

Mutations that affect the PY-NLS or the biophysical properties of its cargo can also reduce Kapβ2 function (16, 20). PY-NLS mutations can be particularly devastating and cause juvenile ALS, as with FUS^P525L^, where the conserved proline of the PY-NLS is mutated to leucine (77, 78). Such aggressive pathology is explained at the molecular level, as Kapβ2 can have significantly lower affinity for PY-NLS mutant protein (14,16,17,23,79). Kapβ2 does show activity against PY-NLS mutants, but higher Kapβ2 concentrations are required for robust effects (16,19,23). Moreover, mutations outside of the PY-NLS can hamper Kapβ2 activity. For example, when Kapβ2 is added to self-assembled FUS harboring specific disease-linked mutations outside the PY-NLS that reduce the dynamics of FUS self-assembly, it can only partially restore wild-type (WT) FUS-like behavior (20, 24). Thus, even when a cargo PY-NLS is intact, other more distributed secondary interactions are critical for Kapβ2 chaperone activity (16, 19). Beyond these insights, however, it remains unclear how Kapβ2 chaperones and disaggregates its cargo.

Here, to probe the mechanism by which Kapβ2 chaperones cargo, we rationally design truncated variants of Kapβ2 and assess their activity. Specifically, we assess their ability to: (1) antagonize FUS aggregation and toxicity in yeast, (2) antagonize FUS condensation at the pure protein level, and (3) antagonize FUS condensation in human cells. We find that a C-terminal fragment of Kapβ2 that contains all cargo-binding residues can perform chaperone activity *in vitro*, and can chaperone FUS in yeast and human cells to a similar extent as Kapβ2. At a minimum, our findings illuminate key mechanistic aspects of Kapβ2 chaperone activity with respect to NIR architecture. Moreover, we reveal a minimal Kapβ2 construct that might be more readily delivered in adeno-associated viruses (AAVs) as a potential therapeutic for several debilitating degenerative disorders.

## Results

### Designing Truncated Kapβ2 Variants to Define the Molecular Determinants of Chaperone Activity

Kapβ2 combines cargo recognition and nuclear delivery in a single NIR (6, 80). Kapβ2 is a superhelical protein that is comprised of 20 HEAT (Huntington, Elongation factor 3, A subunit of protein phosphatase 2A, and Tor1 kinase) repeats (81). Each HEAT repeat is a protein tandem repeat structural motif (∼30–40 residues) composed of two anti-parallel alpha helices (referred to as A- and B-helices) that are typically connected by a short loop (82). The superhelical conformation of Kapβ2 is found in the unbound state, and when Kapβ2 is bound to Ran-GTP or FUS PY-NLS cargo (14,81,83) (**Figure 1A**). Comparing the crystal structure of unbound Kapβ2 with that of Ran-GTP-bound and cargo (FUS PY-NLS)-bound Kapβ2 reveals three functionally distinct segments (84). In the presence of Ran-GTP, an N-terminal segment (N1) consisting of HEAT repeats 1-8 (H1-H8) interacts with Ran-GTP in the nucleus to facilitate cargo release (84) (**Figure 1A**). In the cytoplasm, two C-terminal segments, H9-H13 (C1) and H14-H18 (C2), rotate about a flexible hinge in H13-H14 in response to cargo loading (7,81,84) (**Figure 1A**). In the unbound state, N-terminal H1-H4 and C-terminal H19-H20 exhibit flexibility (84).

**Figure 1.**
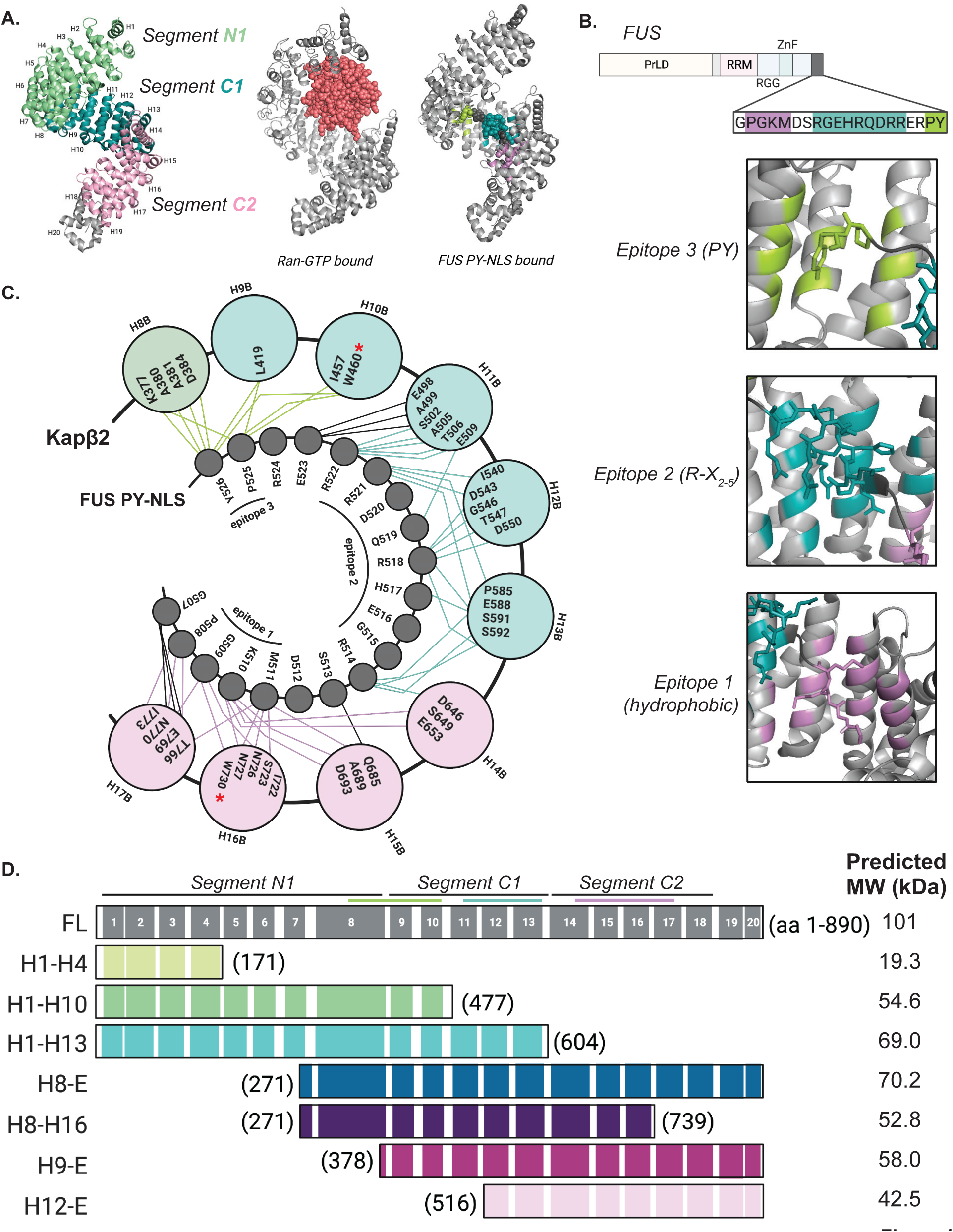
Binding of Kapβ2 to the PY-NLS of FUS is segmental. (**A**) Left, a crystal structure of Kapβ2 in its unbound state (PDB: 2QMR) with each HEAT repeat indicated. Each structural segment is colored: N-terminal segment, N1 (green), the first C-terminal segment, C1 (teal), and the second C-terminal segment, C2 (light pink). Middle, a structure of Kapβ2 in its Ran-GTP bound state, where Kapβ2 is gray and Ran-GTP is salmon (PDB: 1QBK). Right, Kapβ2 is shown bound to the PY-NLS of FUS, where epitope 1 is lilac, epitope 2 is teal, and epitope 3 is lime (PDB: 4FDD). Kapβ2 adopts a largely similar structure in all three states, with slight variations at the N- and C-terminal segments to accommodate binding partners. (**B**) Top, the domain architecture of FUS, with the PY-NLS sequence highlighted. FUS harbors an N-terminal prion-like domain (PrLD), followed by an RNA-recognition motif (RRM), an Arg-Gly-Gly-rich (RGG) domain, a zinc finger (ZnF), and a second RGG domain. Bottom, zoomed-in images of each epitope of the FUS PY-NLS bound to Kapβ2. The residues of Kapβ2 that contact the PY-NLS are colored corresponding to which epitope they interact with, where epitope 3 contacts are colored in lime, epitope 2 contacts are teal, and epitope 1 contacts are lilac. (**C**) A cartoon depiction of Kapβ2 bound to the PY-NLS of FUS, where the outer circles represent each interior (B) helix of the indicated HEAT repeat and the inner circles represent the residues of the FUS PY-NLS. The outer circles are color-coded according by which segment they belong to, with segment N1 shown in green, segment C1 shown in teal, and segment C2 shown in pink. Interactions between Kapβ2 and FUS are shown as lines connecting Kapβ2 residues and FUS residues. Lines are colored to indicate residues making epitope contacts, where epitope 3 contacts are colored in lime, epitope 2 contacts are teal, and epitope 1 contacts are lilac. When Kapβ2 binds to FUS and other cargo, stereotypic residues within the inner helix of each HEAT repeat contact distinct cargo epitopes. W460 and W730, which are mutated to alanine in Kapβ2^W460A:W730A^ are indicated with red asterisks. (**D**) Top, a diagram of Kapβ2 with specific structural features highlighted. The black lines indicate the HEAT repeats (shown as numbered gray boxes) comprising the N1, C1, and C2 segments; the lime, teal, and lilac lines indicate the regions of Kapβ2 that interact with epitope 3, 2, and 1, respectively. Bottom, a diagram of each truncation investigated here, named for the HEAT repeats contained in the construct including the amino acids (aa) contained in each construct, and the predicted molecular weight (MW).

Each segment of Kapβ2 contains residues that engage PY-NLS epitopes (1, 84). H8 of N1 and H9-H10 of C1 interact with the PY residues of epitope 3, H11-H13 of C1 and H14 of C2 interact with epitope 2, and H14-H17 of C2 interact with the hydrophobic/basic epitope 1 (1,14,84) (**Figure 1B, C**). The energetic importance of each epitope of the PY-NLS for Kapβ2 binding varies across different cargo, and so the extent to which Kapβ2 depends on the recognition of any specific epitope for a given cargo protein can be difficult to assess *a priori* (1,12,84,85). However, there are key interactions that appear critical for all cargo, such as the interactions with epitopes 1 and 3 that are impacted by the Kapβ2^W460A:W730A^ mutant (1, 16).

To determine the role of each of the larger structural elements of Kapβ2 with respect to chaperone activity, we designed seven truncated Kapβ2 constructs that span the length of the protein (**Figure 1D**). The truncations were designed to probe the importance of the distribution of Kapβ2 residues that contact cargo, paying special attention to the residues that make direct contact with the PY epitope of the PY-NLS (1,12,14,85).

First, H1-H4 (residues 1-171) was designed to test the N-terminal region of Kapβ2 (**Figure 1D**). This region engages Ran-GTP, but is not known to engage FUS (**Figure 1A**). Hence, we do not anticipate this construct will have chaperone activity.

Second, H1-H10 (residues 1-477) contains segment N1 and the two N-terminal HEAT repeats of segment C1, which contain all the residues that interact with the PY-epitope of FUS, including W460 (**Figure 1C, D**). If H1-H10 can chaperone FUS as effectively as Kapβ2, then recognition of the PY-epitope alone would be sufficient for chaperone activity.

Third, H1-H13 (residues 1-604) contains segments N1 and C1, and contains the residues of Kapβ2 that engage epitopes 2 and 3 (**Figure 1C, D**). If H1-H13 can chaperone FUS as effectively as Kapβ2, then recognition of epitopes 2 and 3 would be sufficient for chaperone activity.

Fourth, H8-E (HEAT repeat 8 to end; residues 271-890) harbors HEAT repeat 8 from N1 plus both C-terminal segments, and contains all the residues that contact each epitope of the FUS PY-NLS (**Figure 1C, D**). If H8-E can chaperone FUS as effectively as Kapβ2, then recognition of epitopes 1, 2, and 3 would be sufficient for chaperone activity.

Fifth, H8-H16 (residues 271-739) harbors a subset of the residues that interact with epitope 1 and all the residues that contact epitope 2 and 3, including W730 (**Figure 1C, D**). If H8-H16 can chaperone FUS as effectively as Kapβ2, then partial recognition of epitope 1 plus contacts with epitope 2 and 3 would be sufficient for chaperone activity.

Sixth, H9-E (residues 378-890) contains segment C1 and C2, and contacts both residues of the PY-epitope, albeit less extensively than H8-E, and contains all contacts for epitopes 1 and 2 (**Figure 1C, D**). If H9-E can chaperone FUS as effectively as Kapβ2, then partial recognition of epitope 3 plus contacts with epitope 1 and 2 would be sufficient for chaperone activity.

Seventh, H12-E (residues 516-890) contains part of segment C1 and all of segment C2, thereby retaining all the contacts for epitope 1 and a subset of contacts for epitope 2 (**Figure 1C, D**). If H12-E can chaperone FUS as effectively as Kapβ2, then the PY-epitope would be dispensable for Kapβ2 to chaperone cargo. We do not expect this possibility given the importance of the PY-epitope integrity for human health (16,23,41,47,77,78). Collectively, this set of Kapβ2 truncations will enable us to define structural and sequence characteristics of Kapβ2 that are critical for its chaperone activity.

### A C-terminal Fragment of Kapβ2, H8-E, Mitigates FUS Toxicity in Yeast

To determine whether truncated Kapβ2 proteins retain chaperone activity, we expressed each construct in a yeast model of FUS proteinopathy. For these studies, we utilized a yeast strain lacking Hsp104, a powerful protein disaggregase that is not found in humans (26,86–88). Overexpression of FUS in this system elicits cytoplasmic FUS aggregation and toxicity, thereby mimicking phenotypes of degenerating neurons in ALS/FTD with FUS pathology (35,89–91). Indeed, elevated FUS expression is linked to ALS (92, 93). Our yeast model of FUS proteinopathy has illuminated several genetic modifiers of FUS aggregation and toxicity, which have translated to fly, mammalian cell and neuronal models of ALS/FTD (16,18,89–91,94–99). Additionally, the yeast homolog of Kapβ2, Kap104, does not engage FUS effectively and cannot transport FUS into the nucleus, providing a simple system for measuring the specific activity of Kapβ2 (13,16,90).

First, we tested whether expression of the Kapβ2 constructs on their own was toxic in yeast. Generally, the Kapβ2 constructs were not overtly toxic (**Figure S1A**). Unexpectedly, however, H1-H10 exhibited some toxicity as did H8-E to a lesser extent (**Figure S1A**). Next, we tested whether truncated forms of Kapβ2 could mitigate FUS toxicity in yeast. Full-length (FL), WT Kapβ2 mitigates FUS toxicity in yeast (**Figure 2A**) without affecting FUS expression level (**Figure 2B**). Of the Kapβ2 truncations tested here, those which lack the complete set of FUS-interacting residues (**Figure 1C,D**), did not mitigate FUS toxicity (**Figure 2A**). Thus, H1-H4 (no FUS-binding residues), H1-H10 (lacks epitope 1 and 2-binding residues), H1-H13 (lacks epitope 1-binding residues), H8-H16 (lacks epitope 1-binding residues in H17), H9-E (lacks epitope 3-binding residues in H8), and H12-E (lacks epitope 2 and 3-binding residues in H8-H11) did not mitigate FUS toxicity (**Figure 1C, 2A**). Perhaps most surprising was the ineffectiveness of H9-E and H8-16, each of which only lack a few FUS-binding residues and retain both W460 and W730 (**Figure 1C**). Thus, the absence of even a few FUS-binding residues reduces the ability of Kapβ2 to mitigate FUS toxicity in yeast.

**Figure 2.**
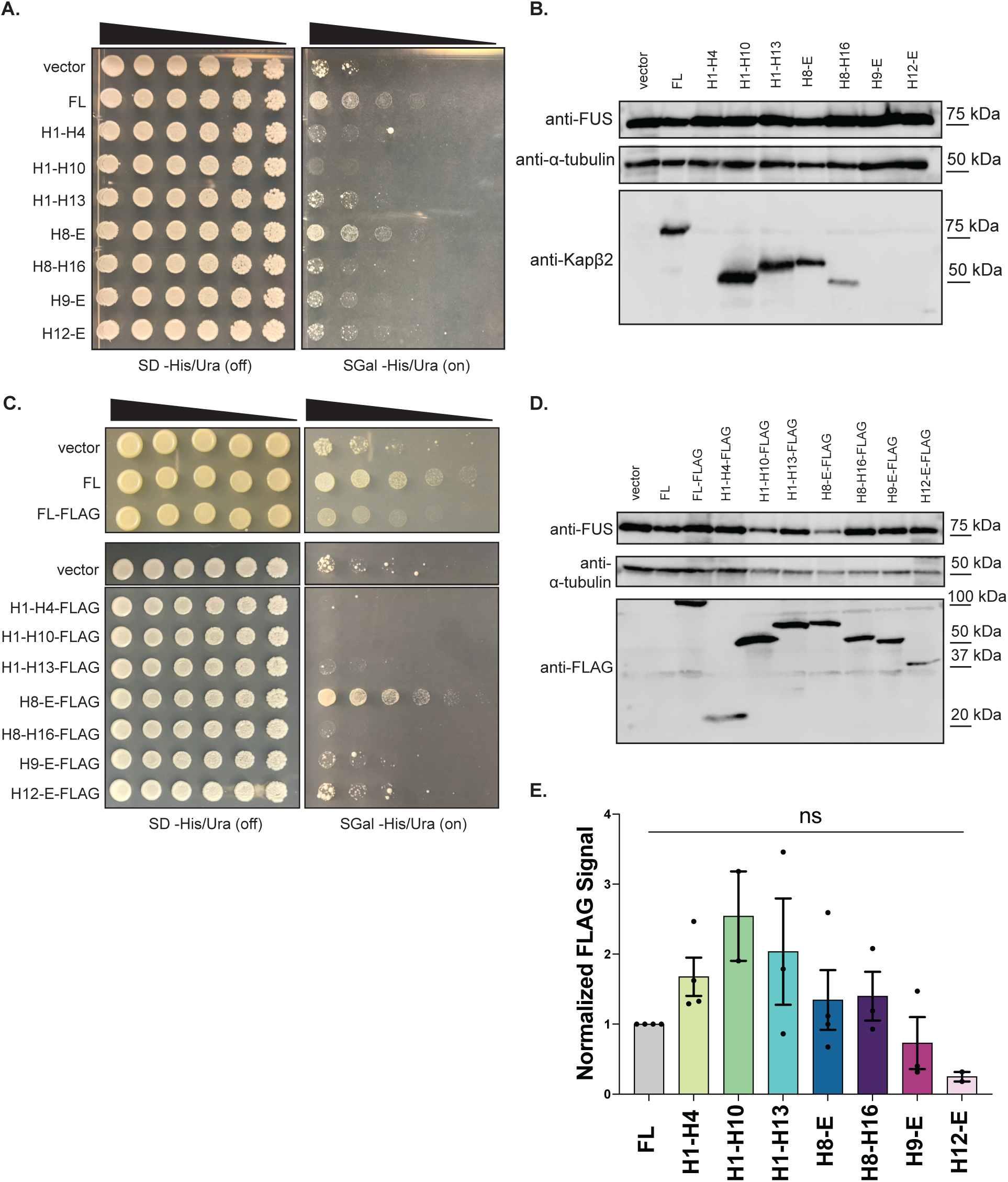
Kapβ2 and H8-E mitigate FUS toxicity in yeast. (**A**) Δ*hsp104* yeast with galactose-inducible FUS integrated were transformed with the indicated galactose-inducible Kapβ2 construct and serially diluted 3-fold onto agar plates with either glucose (expression off) or galactose (expression on) as the carbon source. In the absence of Kapβ2 (vector), FUS expression is very toxic. This toxicity can be mitigated by the expression of full-length (FL) Kapβ2. Growth is also restored with the expression of a C-terminal fragment, H8-E. None of the other Kapβ2 truncations mitigate FUS toxicity. (**B**) Western blot showing FUS and Kapβ2 expression for each condition. The Kapβ2 antibody recognizes residues 298-389 in the eighth HEAT repeat, and thus H1-4, H9-E, and H12-E are not recognized. However, FUS expression is equal in all conditions as compared to the loading control, α-tubulin. (**C**) FLAG-tagged Kapβ2 constructs were transformed into Δ*hsp104* yeast with integrated galactose-inducible FUS, and serially diluted 3-fold onto agar plates with either glucose or galactose as the carbon source. FL Kapβ2-FLAG and H8-E-FLAG suppress FUS toxicity, whereas other Kapβ2 truncations do not. (**D**) A western blot illustrates that most truncations are expressed to similar levels, with H9-E and H12-E showing slightly lower expression relative to the loading control, α-tubulin. **(E)** Western blots were quantified by first normalizing the intensity of the FLAG band to the intensity of the loading control, α-tubulin. The ratio for FL Kapβ2 was then set as 1, and all other values were normalized to FL. Data points represent individual blots, bars represent means ± SEM, n=2-4. An ordinary one-way ANOVA with Dunnett’s multiple comparisons test was used to compare the normalized intensity of the FLAG band for each condition to FL Kapβ2-FLAG.

We also tested Kapβ2^W460A:W730A^ (**Figure S1B-D**), which has just two FUS-binding residues mutated. Kapβ2^W460A:W730A^ was slightly toxic when expressed alone in yeast (**Figure S1D**). Kapβ2^W460A:W730A^ was unable to effectively mitigate FUS toxicity, despite being expressed at similar levels to Kapβ2 (**Figure S1B,C**). Thus, mutating just two residues in Kapβ2 that engage epitope 1 (W730) and epitope 3 (W460) of the FUS-PY-NLS, reduces Kapβ2 activity (16).

In contrast, an N-terminal Kapβ2 truncation that begins at H8 (H8-E, residues 271-890; **Figure 1D**) mitigates FUS toxicity without affecting FUS expression level (**Figure 2A,B**). H8-E is the only truncation tested here that contains all the Kapβ2 residues that bind the FUS PY-NLS, as well as the acidic loop in H8, which is necessary for cargo release in the nucleus (1,12,14). Our findings suggest that the entire cargo-binding surface of Kapβ2 is required for Kapβ2 to mitigate FUS toxicity in yeast.

There are several possible explanations for why an N- or C-terminally truncated Kapβ2 construct might not mitigate FUS toxicity despite retaining the residues known to contact each epitope of the PY-NLS, including the critical Trp residues at positions 460 and 730. One possible explanation is that a truncation construct is not stable or is expressed poorly. However, low expression did not appear to be an issue. Using an antibody that recognizes an epitope in H8 of Kapβ2, we found that H1-H10, H1-H13, H8-E, and H8-H16 were all expressed in yeast, although H8-H16 was expressed at lower levels (**Figure 2B**). We were unable to detect H1-H4, H9-E, and H12-E with this antibody (**Figure 2B**). Thus, to detect the other Kapβ2 truncations that lack H8, we added a C-terminal FLAG tag to each construct to enable detection via immunoblot using anti-FLAG antibodies.

Next, we tested the toxicity of the FLAG-tagged Kapβ2 truncations in the absence of FUS expression. Generally, the Kapβ2-FLAG constructs were not toxic, although Kapβ2-FLAG and H8-E-FLAG were very slightly toxic, whereas H1-H10-FLAG was toxic (**Figure S1E**). As with untagged Kapβ2 constructs, only FL Kapβ2-FLAG and H8-E-FLAG mitigated FUS toxicity in yeast without affecting FUS expression (**Figure 2C, D**). All other Kapβ2 truncations were ineffective (**Figure 2C**). The activity of FL Kapβ2-FLAG was reduced compared to untagged FL Kapβ2 despite similar expression levels, but Kapβ2-FLAG could still mitigate FUS toxicity (**Figure 2C, S1F**). H8-E-FLAG strongly reduced FUS toxicity (**Figure 2C**). Western blots revealed that although there was some variation in Kapβ2-FLAG construct expression, these differences were not statistically significant (**Figure 2D, E**). Thus, all the Kapβ2 constructs were robustly expressed. However, only FL Kapβ2 and H8-E mitigate FUS toxicity. Our findings suggest that when any single set of epitope contacts is lost, Kapβ2 is no longer able to mitigate FUS toxicity in yeast.

### Truncated Kapβ2 Proteins Promote FUS Solubility and Nuclear Localization in Yeast

We next determined whether truncated Kapβ2 variants could suppress FUS aggregation and promote nuclear FUS localization in yeast. We did not explore the H1-H10 construct further due to its toxicity (**Figure S1A,E**). Yeast cells expressing FUS-GFP form cytoplasmic aggregates (89-91,96). Previously, we have found that potentiated variants of the Hsp104 disaggregase can disperse these FUS aggregates without localizing FUS to the nucleus (91,94–96). By contrast, FL Kapβ2 suppresses FUS aggregation in yeast and localizes FUS to the nucleus (**Figure 3A-C**) (16). For the Kapβ2 truncations expressed here, the ratio of FUS-GFP expression to the loading control, α-tubulin, is relatively constant (**Figure 3D**). Thus, the chaperone activity observed is attributable to Kapβ2 activity, and not degradation or low expression of FUS.

**Figure 3.**
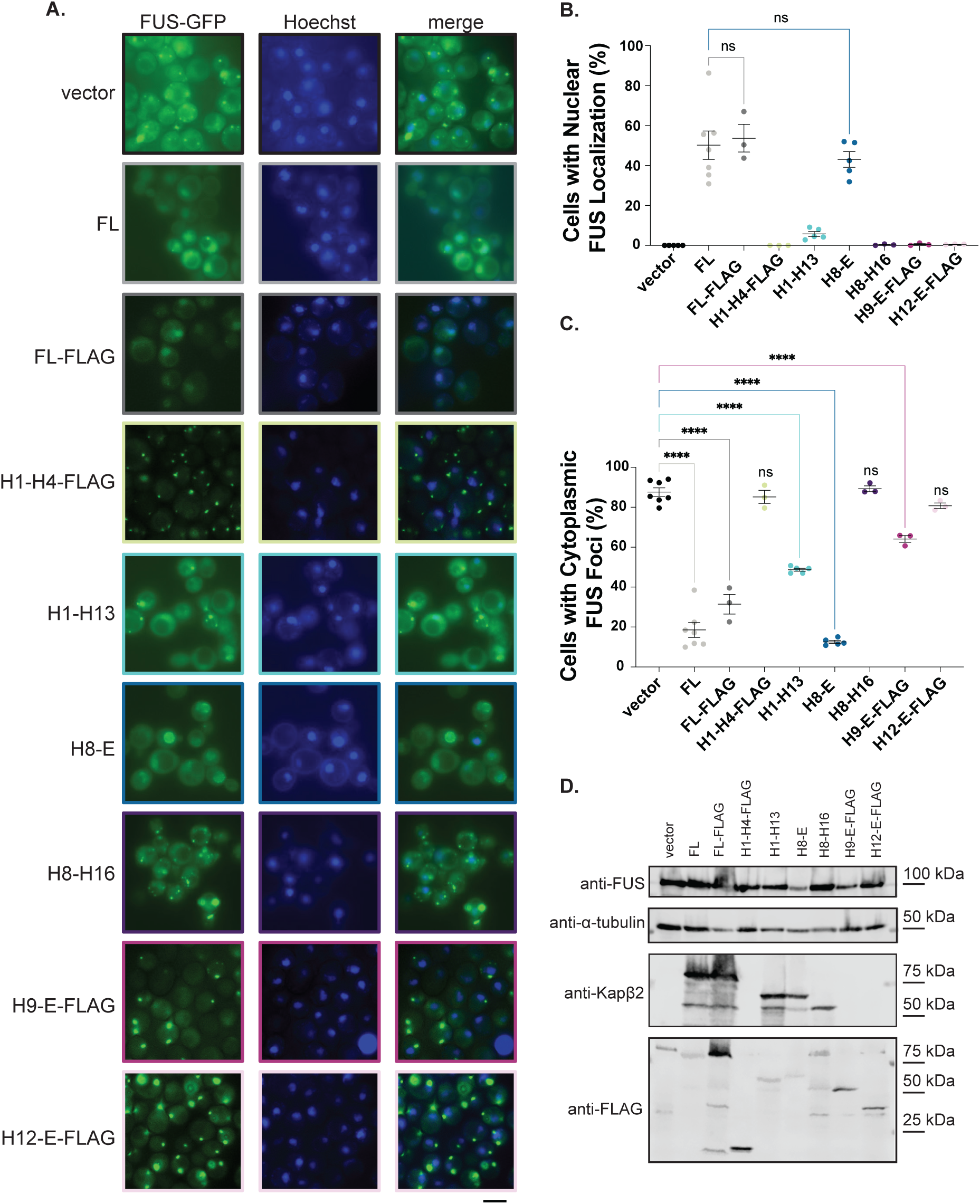
Kapβ2 and H8-E resolve cytoplasmic FUS foci and promote nuclear localization of FUS in yeast. (**A**) Δ*hsp104* yeast with integrated galactose-inducible FUS-GFP were transformed with the indicated galactose-inducible Kapβ2 truncation and grown in galactose-containing media for 5 h. At this time, fluorescence microscopy reveals that yeast without Kapβ2 harbor cytoplasmic FUS foci. In contrast, cells expressing FL Kapβ2 (untagged or with a C-terminal FLAG tag) have a low proportion of cells with cytoplasmic FUS foci and a large proportion of cells with nuclear FUS localization. Similar results are observed with H8-E. Expression of H1-H13 or H9-E-FLAG can reduce the proportion of cells with cytoplasmic FUS foci without restoring nuclear FUS localization. H1-H4-FLAG, H8-H16, and H12-E-FLAG are unable to decrease the proportion of cells with cytoplasmic FUS foci or increase the proportion of cells with nuclear FUS localization. Scale bar = 5 µm. (**B**) Quantification of the percentage of cells with nuclear FUS localization for each condition. Points represent independent transformations, and at least 100 cells were counted per transformation. Values represent means ± SEM, n=3-7. For cells with nuclear FUS localization, an ordinary one-way ANOVA with Dunnett’s multiple comparisons test was used to compare the percentage of cells with nuclear localization in each condition to cells expressing FL Kapβ2. (**C**) Quantification of the percentage of cells with cytoplasmic FUS foci. Points represent independent transformations, and at least 100 cells were counted per transformation. Values represent means ± SEM, n=3-7. An ordinary one-way ANOVA with Dunnett’s multiple comparisons test was used to compare the percentage of cells with cytoplasmic FUS foci in each condition to the vector control; ****, p<0.0001. (**D**) Western blot of FUS-GFP yeast transformed with vector alone, FL Kapβ2, or the indicated Kapβ2 truncation. Probing with anti-FUS and anti-alpha-tubulin shows similar levels of protein expression across samples. Kapβ2 truncation expression, however, does vary, with H9-E-FLAG and H12-E-FLAG expressed at low levels relative to FL Kapβ2-FLAG.

Among the truncated Kapβ2 proteins, we observed a range of phenotypes. H1-H4-FLAG, H8-H16, and H12-E-FLAG were unable to prevent FUS aggregation or localize it to the nucleus (**Figure 3A-C**). By contrast, H1-H13 and H9-E-FLAG exhibited an intermediate phenotype. These truncations were unable to localize FUS to the nucleus, but did reduce the percentage of cells with cytoplasmic foci (**Figure 3A-C**). This phenotype is distinct from yeast expressing FUS-GFP and Kapβ2^W460A:W730A^, which neither dissolves cytoplasmic FUS structures nor promotes FUS nuclear localization (16). Western blot confirmed that the lower level of cytoplasmic FUS foci is not due to increased expression of the Kapβ2 truncation, as H9-E-FLAG is expressed at lower levels than Kapβ2-FLAG (**Figure 3D**). Thus, a decrease in cytoplasmic FUS foci without increased nuclear localization suggests that, in yeast, H1-H13 and H9-E-FLAG may partially inhibit FUS self-assembly in the cytoplasm, but are inactive as NIRs.

Furthermore, this partial inhibition of FUS aggregation by H1-H13 and H9-E-FLAG was insufficient to mitigate FUS toxicity (**Figure 2A,C**). FUS must aggregate in the cytoplasm *and* bind RNA to confer toxicity in yeast (Sun et al. 2011). Kapβ2 can cause FUS to eject bound RNA (17, 19). Hence, it may be that truncated Kapβ2 proteins have reduced ability to eject bound RNA from FUS. Thus, FUS would continue to sequester important RNA transcripts and confer toxicity in yeast. Alternatively, the limited effect on FUS aggregation by H1-H13 or H9-E-FLAG may be insufficient to reduce FUS toxicity.

Consistent with the ability to mitigate FUS toxicity as well as Kapβ2, the H8-E construct robustly suppresses FUS aggregation and localizes FUS to nucleus (**Figure 3A,B**). This finding suggests that all the information needed to chaperone FUS and transport it to the nucleus is encoded in H8-E.

### A C-terminal Fragment of Kapβ2, H8-E, Robustly Prevents FUS Aggregation in Vitro

For a more direct assessment of the chaperone activity of truncated Kapβ2 proteins, we purified recombinant Kapβ2, Kapβ2^W460A:W730A^, various Kapβ2 truncations (**Figure S2A**), and FUS to measure chaperone activity *in vitro*. FUS is intrinsically aggregation-prone, and thus to maintain solubility during purification, FUS was purified with an N-terminal GST-tag (90). The bulky nature of GST precludes the PrLDs of FUS molecules from interacting with each other, thus preventing fibrillization (90). Additionally, there is a linker between the GST-tag and FUS which contains a TEV protease cleavage site. Thus, TEV protease can be utilized to initiate FUS fibrillization, which can be tracked via turbidity measurements (16, 90). The FUS fibrils that form *in vitro* resemble the fibrils observed in postmortem tissue and are toxic to neurons in culture, and thus represent a biologically relevant structure to study (61, 90).

Purification of Kapβ2, Kapβ2^W460A:W730A^, H1-H13, and H8-E was routine (**Figure S2A**). However, we noticed that the expression, yield, and purity of H9-E were lower than for H1-H13 or H8-E (**Figure S2A**). Previous work has indicated that HEAT-repeat proteins can be sensitive to truncation (100, 101). In structural studies of other HEAT-repeat proteins, some truncations are stable, whereas others are aggregation-prone (100, 101). Indeed, purified H8-H16, which is inactive in yeast (**Figure 2A, C, 3A-C**), is even more unstable than H9-E *in vitro* and becomes insoluble upon cleavage of its His-SUMO solubility tag by ubiquitin-like-specific protease 1 (Ulp1), and thus was purified without cleavage (**Figure S2A**). FL Kapβ2 chaperone activity is nearly identical between the Ulp1-cleaved and uncleaved constructs (**Figure S2B, C**), and thus we can rule out any effect of the His-SUMO tag itself on chaperone activity for His-SUMO-H8-H16. The instability of H8-H16 suggests that C-terminal H17-H20 are important for protein stability in the absence of H1-H7.

Next, we assessed the extent of FUS aggregation in the presence of increasing concentrations of each Kapβ2 construct (**Figure 4A-F, S2D, E**). FL Kapβ2 suppresses FUS aggregation at sub-stoichiometric levels (**Figure 4A, S2E**) (16). By contrast, Kapβ2^W460A:W730A^ does not effectively prevent FUS aggregation, even at over 3-fold excess (**Figure S2D,E**). Likewise, purified His-SUMO-H8-H16 and H9-E were unable to effectively prevent FUS aggregation (**Figure 4E,F, S2E**). In agreement with the reduction in cytoplasmic FUS-GFP foci in yeast expressing H1-H13, H1-H13 can partially inhibit FUS aggregation *in vitro*, but only at very high levels (**Figure 4B, S2E**). Indeed, at the highest concentration tested, H1-H13 reduces FUS aggregation to only ∼40% (**Figure 4B, S2E**). In line with the suppression of FUS aggregation and toxicity observed in yeast, H8-E robustly prevents aggregation of FUS *in vitro* (**Figure 4C, S2E**). Calculating the half maximal inhibitory concentration (IC_50_) of each construct for prevention of FUS aggregation reveals that H8-E can prevent FUS aggregation as effectively as FL Kapβ2 (**Figure 4D**). Our findings suggest that the entire FUS-binding region of Kapβ2 is required to enable maximal inhibition of FUS aggregation *in vitro*.

**Figure 4.**
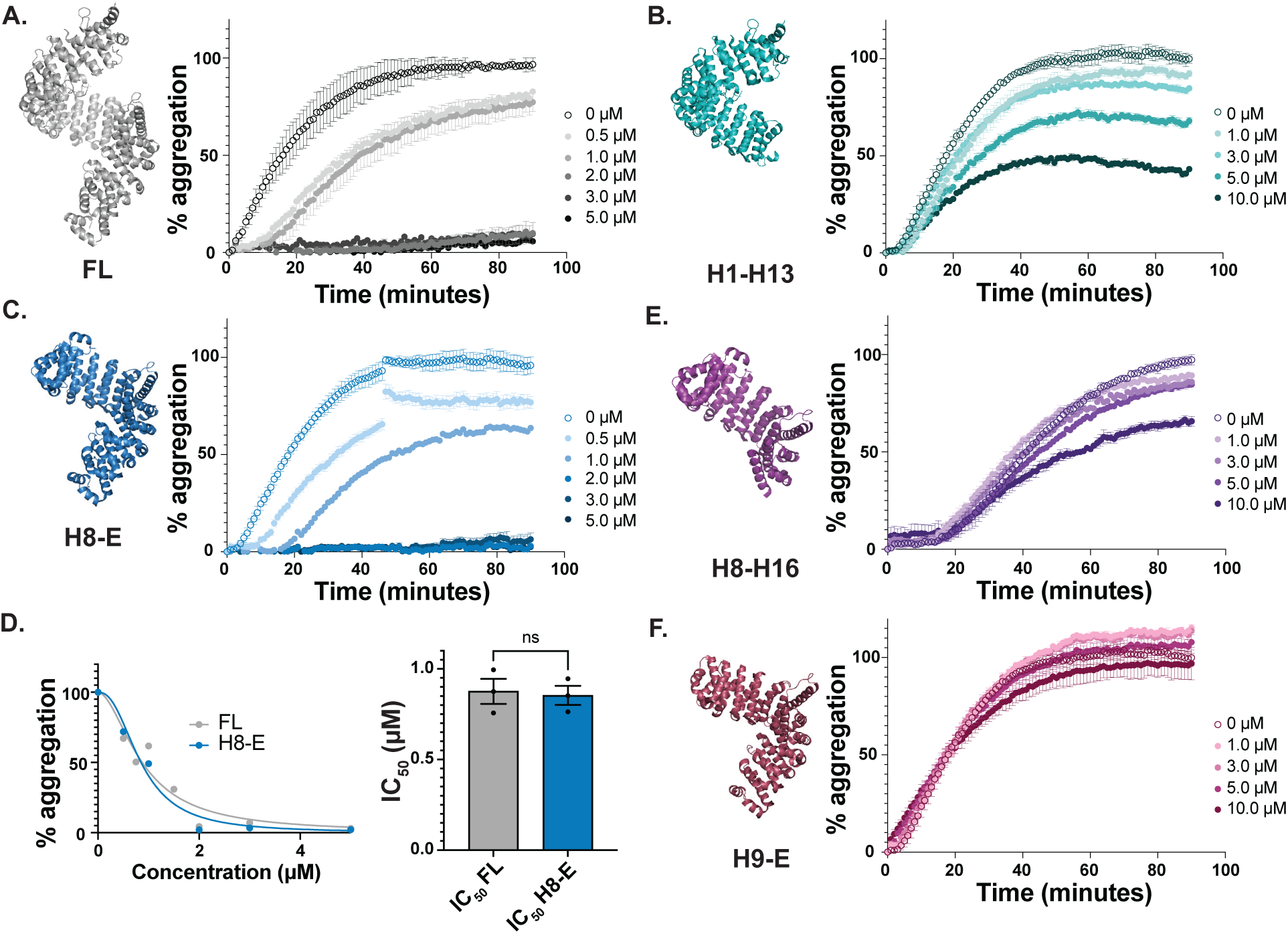
Kapβ2 and H8-E robustly suppress FUS aggregation. To measure the ability of Kapβ2 constructs to prevent FUS aggregation, we incubated 3 μM FUS with 1 μg TEV protease and the indicated concentration of Kapβ2 and monitored turbidity at 395 nm. Increased turbidity indicates FUS self-assembly and each curve represents the mean of 2-4 trials ± SEM. For each experiment, the maximal value of FUS aggregation in the absence of Kapβ2 was set to 100 and all data for that trial were normalized to that value. (**A-C**) Kinetics of FUS aggregation in the presence of increasing concentrations of Kapβ2 (A), H1-13 (B), or H8-E (C). FL Kapβ2 strongly suppresses FUS aggregation. H1-13 modestly inhibits FUS aggregation, but only at stoichiometric excess. H8-E strongly suppresses FUS aggregation. (**D**) To compare the ability of FL Kapβ2 and H8-E to suppress FUS aggregation, the area under the curve produced at each concentration was used to construct a curve of aggregation vs. concentration (left). Comparing the half maximal inhibitory concentration (IC_50_) reveals that the ability of H8-E to prevent FUS aggregation is very similar to FL Kapβ2. IC_50_ values were derived from a non-linear curve fit with normalized responses and a variable slope. Each calculated IC_50_ value is plotted with its 95% confidence interval. An unpaired t-test was used to compare IC_50_ values. (**E, F**) Kinetics of FUS aggregation in the presence of increasing concentrations of His-SUMO-H8-H16 (E) or H9-E (F). High concentrations of His-SUMO-H8-H16 slightly inhibit FUS aggregation. H9-E is unable to prevent FUS aggregation even in excess. The structures presented here are based on PDB: 2QMR.

### H8-E Reverses FUS Aggregation in Vitro Nearly as Effectively as Kapβ2

In addition to preventing the aggregation of its cargo, Kapβ2 can also reverse the formation of pre-formed fibrils (16). However, more Kapβ2 is required to reverse FUS aggregation than to prevent aggregation (16). Thus, focusing on the truncations that had the most significant effect on preventing FUS aggregation, H1-H13 and H8-E, we next determined whether truncated variants of Kapβ2 could disaggregate pre-formed FUS fibrils. For these experiments, FUS aggregation was initiated by the addition of TEV protease in the absence of Kapβ2. After one hour of FUS aggregation when abundant FUS fibrils are present (90), Kapβ2 was added, and turbidity was monitored.

As expected, FL Kapβ2 rapidly disaggregates FUS fibrils, completely dissolving FUS aggregates after 10 minutes at a ratio of ∼3:1 Kapβ2 to FUS (**Figure 5A**) (16, 102). By contrast, Kapβ2^W460A:W730A^ was unable to dissociate preformed FUS fibrils (**Figure S3A, B**). H1-H13 was also ineffective at disaggregating FUS, but displayed some activity (**Figure 5B, S3B**). At the highest concentration tested, H1-13 was able to reduce turbidity by ∼50% after 80 minutes (**Figure 5B**). Additionally, the shape of the FUS disaggregation curve for H1-H13 is distinct from what is observed with FL Kapβ2 (**Figure 5A, B)**. For FL Kapβ2, FUS disaggregation begins immediately after addition and decreases rapidly (**Figure 5A**). By contrast, addition of H1-H13 initially causes a sharp reduction in FUS aggregation, followed by apparent re-aggregation and subsequent slow disaggregation (**Figure 5B**). These data indicate that H1-H13 has limited ability to effectively disaggregate FUS.

**Figure 5.**
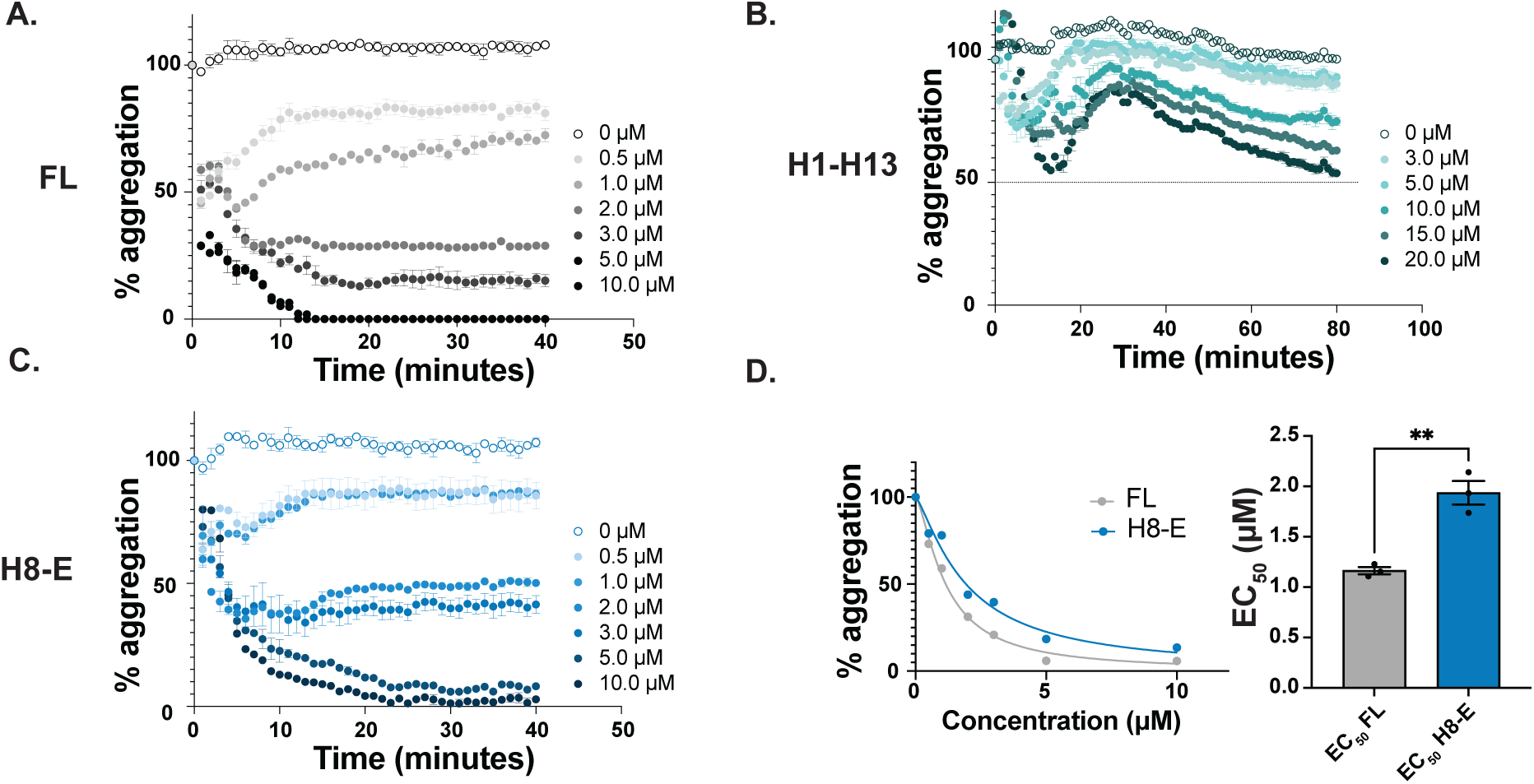
Kapβ2 and H8-E robustly reverse FUS aggregation. To measure the ability of Kapβ2 constructs to reverse FUS aggregation, we incubated 3 μM FUS with 1 μg TEV protease and monitored turbidity at 395 nm. After one hour of aggregation, the indicated amount of Kapβ2 was added and turbidity was continually measured. Each curve represents the mean of 2-3 trials ± SEM. For each condition, the value of FUS aggregation immediately prior to adding Kapβ2 was set to 100 and all subsequent readings were normalized to that value. (**A-C**) Kinetics of FUS disaggregation by FL Kapβ2 (A), H1-H13 (B), or H8-E (C). FL Kapβ2 and H8-E rapidly disaggregate FUS. H1-H13 can partially reverse FUS aggregation, but only to ∼50% and over an extended period. (**D**) To compare activity of FL Kapβ2 and H8-E, the area under the curve produced at each concentration was used to construct a curve of aggregation vs. concentration (left). Half maximal effective concentration (EC_50_) values were derived from a non-linear curve fit with normalized responses and a variable slope. Each calculated EC_50_ value is plotted with its 95% confidence interval. Comparing the EC_50_ reveals that the ability for H8-E to reverse FUS aggregation is somewhat impaired relative to Kapβ2. An unpaired t-test was used to compare EC_50_ values. **, p<0.005.

Importantly, H8-E is a robust chaperone which readily disperses pre-formed FUS aggregates (**Figure 5C, S3B**). However, H8-E requires a longer time to completely disaggregate FUS, as complete disaggregation is not achieved until ∼20 minutes (**Figure 5C**), whereas FL Kapβ2 only requires ∼10 minutes (**Figure 5A**). Moreover, the half maximal effective concentration (EC_50_) of H8-E required to disaggregate FUS is ∼2-fold higher than for FL Kapβ2 (**Figure 5D**). Thus, H8-E can disaggregate FUS at sub-stoichiometric levels, but it is unable to disaggregate FUS as well as FL Kapβ2. Our findings suggest that the N-terminal H1-H7 region of Kapβ2 is more critical for optimal reversal of FUS aggregation than for inhibition of FUS aggregation. Indeed, the N-terminal H1-H7 region of Kapβ2 appears to enable optimal disaggregation activity.

### H8-E Dissolves Preformed FUS Liquid Condensates

FUS undergoes liquid-liquid phase separation to spontaneously form dynamic condensates, which are important for FUS functionality, but may precede pathological aggregation, and thus require vigilant chaperoning (53,60,64,69,103,104). *In vitro*, FUS readily forms liquid condensates in the presence of Cy3-labeled poly(U) RNA (20). Thus, we next tested if the Kapβ2 truncations that effectively work on solid-like FUS fibrils would also dissolve heterotypic liquid-liquid phase separated FUS:RNA condensates. FL Kapβ2 can dissolve preformed FUS:RNA condensates, resolubilizing both the RNA (**Figure 6A**) and FUS (**Figure 6B**) components of these condensates (20,21,24). FL Kapβ2 reduces the number of FUS:RNA droplets (**Figure 6C**) and the total area of these structures (**Figure 6D**). H1-H13 also significantly reduced the number of FUS:RNA droplets after addition, but only by ∼50% as opposed to the near 100% reduction observed for FL Kapβ2 (**Figure 6A-D**). Moreover, the total area occupied by the remaining droplets is not lower than the condition in which no Kapβ2 was added, indicating that there are fewer, larger droplets present after treatment with H1-H13. Thus, partial recognition of FUS by the H1-H13 truncation variant appears to induce droplet fusion (**Figure 6B**). These data are consistent with the limited ability of H1-H13 to antagonize FUS aggregation (**Figure 5B**).

**Figure 6.**
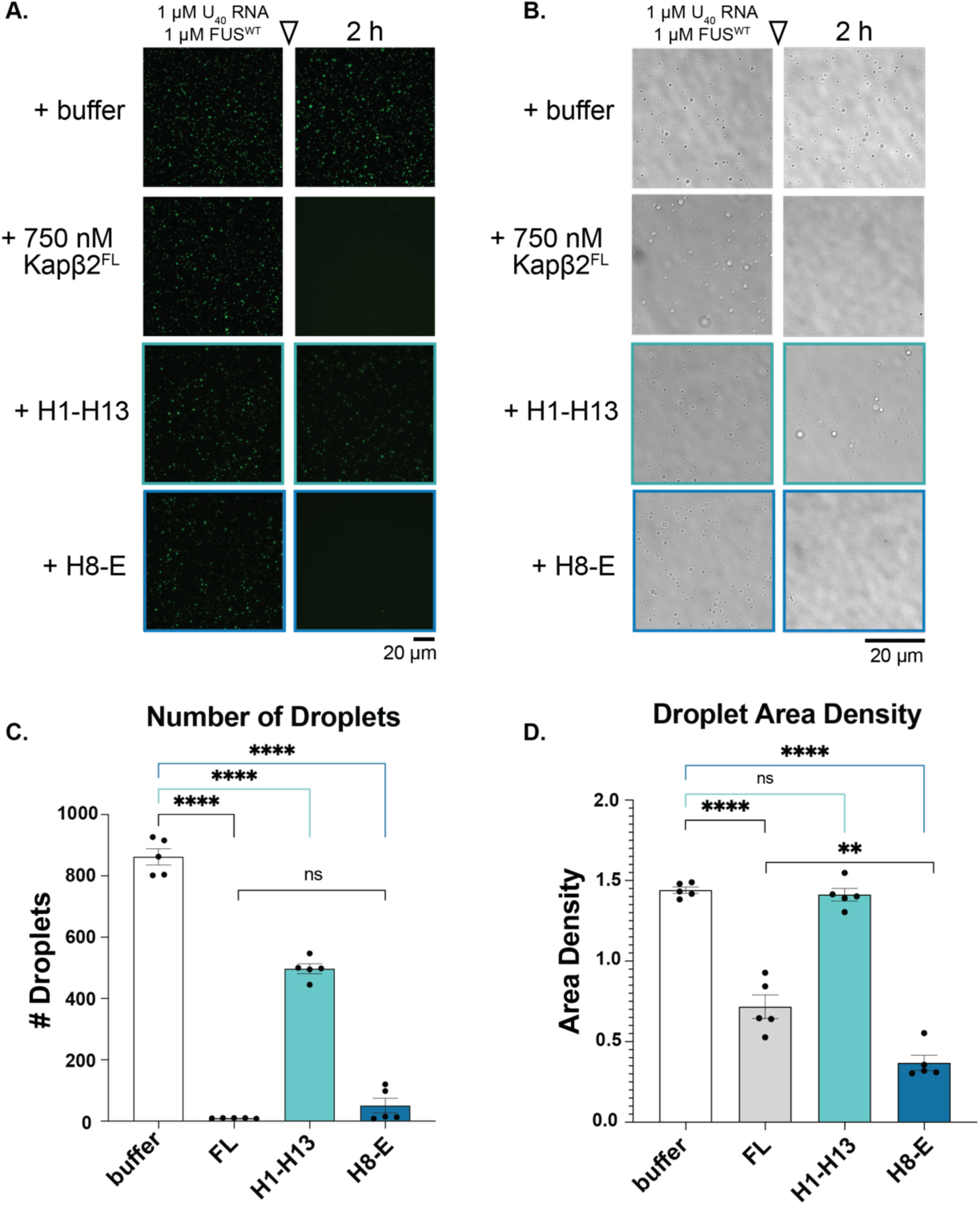
Kapβ2 and H8-E disperse heterotypic FUS:RNA condensates. (A-D) To assess the ability of Kapβ2 and truncation constructs to disperse FUS:RNA droplets, we mixed 1 µM Cy3-labeled U_40_ RNA with 1 µM FUS, which form liquid-like droplets after 4 h. Taking images using fluorescence microscopy (A) or brightfield microscopy (B) 2 h after the addition of buffer or the indicated Kapβ2 construct demonstrates that FUS:RNA droplets are unaffected by the addition of buffer (scale bar = 20 μm). By contrast, FL Kapβ2 (0.75µM) effectively disperses FUS:RNA droplets, reducing them in number (C) and in the overall area of the coverslip that was occupied by condensates (D). H1-H13 can also reduce the number of FUS:RNA droplets, but is not as effective as FL Kapβ2. However, an equal concentration of H8-E can reduce the number and area of FUS:RNA droplets to a similar extent as FL Kapβ2. Data points in (C) and (D) represent individual images, and bars represent mean ± SEM, n = 5. An ordinary one-way ANOVA with Dunnett’s multiple comparisons test was used to compare the cytoplasmic to nuclear GFP-FUS ratio for each condition to that of buffer alone. ****, p<0.0001. Additionally, an unpaired t-test was used to compare FL and H8-E. **, p<0.004.

Like FL Kapβ2, H8-E significantly reduced the number and total area of FUS:RNA droplets, performing as effectively as Kapβ2 (**Figure 6A-D**). The total number of droplets resulting after treatment with Kapβ2 and H8-E is similar (**Figure 6C**), but the total area of the droplets is lower for H8-E (**Figure 6D**). Thus, the few droplets that remain in the presence of H8-E are even smaller than those that remain with Kapβ2. Our findings suggest that H8-E effectively dissolves FUS liquids, and in this context the N-terminal H1-H7 of Kapβ2 is less important. Hence, it appears that the N-terminal H1-H7 region of Kapβ2 is critical for optimal disaggregation of solid FUS condensates but not liquid FUS condensates.

### Truncated Kapβ2 Proteins Reduce Cytoplasmic Foci and Restore Nuclear Localization in Human Cells

Finally, we explored how Kapβ2 and truncation variants might affect FUS condensation and localization in human cells. The overexpression of GFP-FUS plus mCherry (negative control) in Henrietta Lacks’ (HeLa) cells induces the formation of cytoplasmic FUS foci (**Figure 7A,B**). Coexpression of mCherry-FL-Kapβ2 significantly reduced the formation of cytoplasmic FUS foci (**Figure 7A,B**). In contrast, expression of mCherry-Kapβ2^W460A:W730A^ exacerbates formation of cytoplasmic FUS foci (**Figure 7A,B**). Of the truncation constructs, neither mCherry-H1-H10 nor mCherry-H12-E reduced the percentage of cells with cytoplasmic FUS foci (**Figure 7A,B**). Coexpression of mCherry-H8-E or mCherry-H1-H13 reduced the percentage of cells with cytoplasmic FUS foci to a similar extent as Kapβ2 (**Figure 7A,B**). Unexpectedly, H9-E also reduced formation of cytoplasmic FUS foci (**Figure 7A,B**). mCherry-H8-H16 was barely visible via microscopy, and was not detected via western blot, and thus was not analyzed further (**Figure S4**). Moreover, there was variable expression of each expressed Kapβ2 construct (**Figure S4**). FL Kapβ2, Kapβ2^W460A:W730A^, H1-H10, and H12-E were each robustly expressed, whereas H1-H13, H8-E, and H9-E were expressed at lower levels (**Figure S4**). However, despite the low expression, H1-H13, H8-E, and H9-E mitigated formation of cytoplasmic FUS foci (**Figure 7A,B**). Moreover, GFP-FUS was similarly expressed across all cells relative to the α-tubulin loading control (**Figure S4**). Thus, differences in FUS expression do not contribute to the reduced formation of cytoplasmic FUS foci.

**Figure 7.**
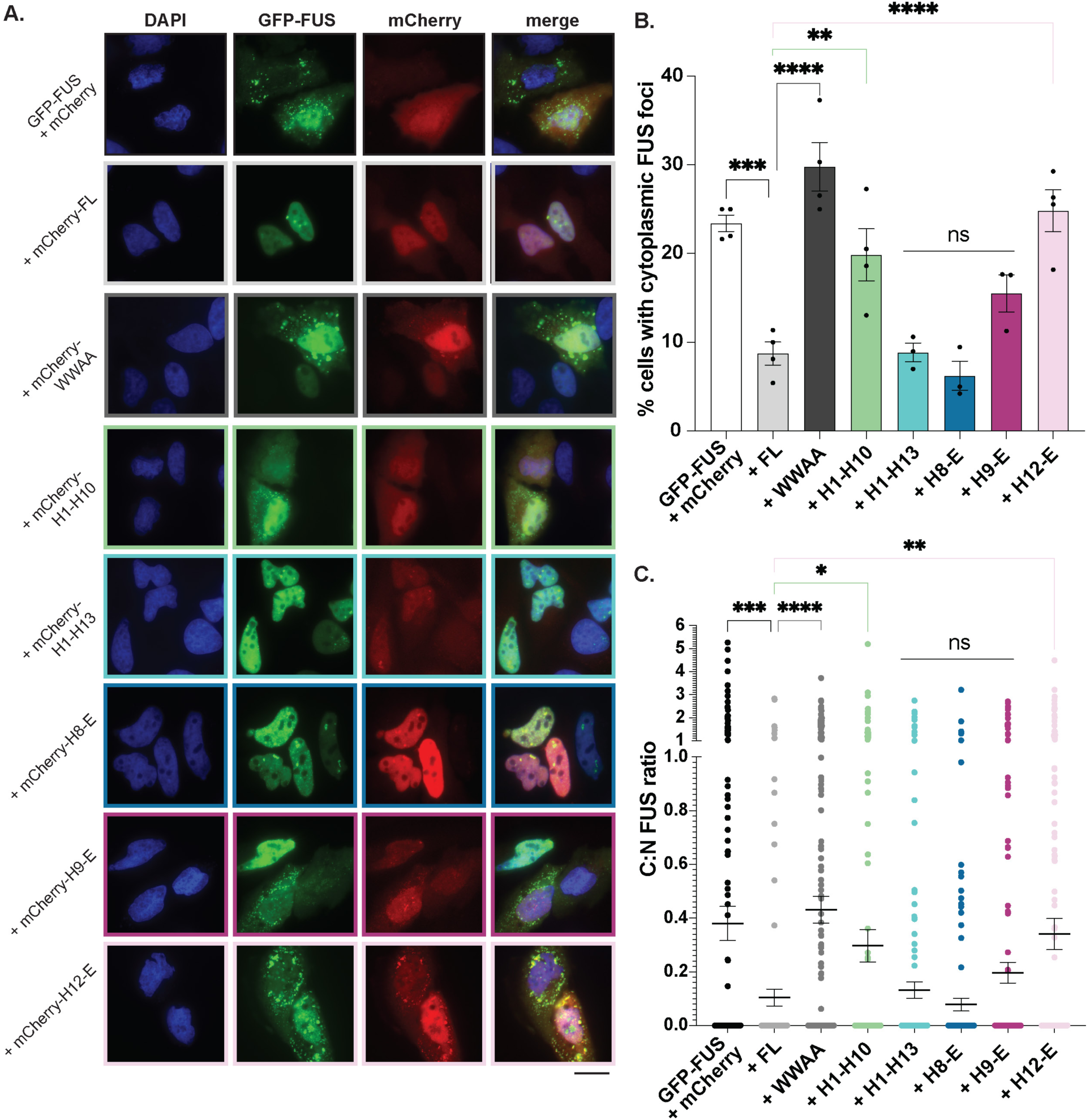
Kapβ2 and H8-E prevent the formation of cytoplasmic FUS foci and promote FUS nuclear localization in human cells. (**A**) In HeLa cells, expression of GFP-FUS with mCherry alone results in ∼25% of cells with cytoplasmic FUS puncta. When mCherry-FL-Kapβ2 is co-expressed with GFP-FUS, FUS is largely localized to the nucleus, with many fewer cells exhibiting cytoplasmic mislocalization. Furthermore, expression of H1-H13, H8-E, or H9-E also reduces the proportion of cells with cytoplasmic FUS foci. In contrast, cytoplasmic foci persist when H1-H10 and H12-E are expressed. These results suggest that in human cells where endogenous FL Kapβ2 is present, H1-H13 and H9-E can complement the endogenous chaperone machinery to antagonize FUS condensation and promote nuclear import. Scale bar = 20 µm. (**B**) Quantification of the percentage of cells with cytoplasmic FUS foci indicates that the activity of each H1-H13, H8-E, and H9-E each is not statistically different from FL Kapβ2. Each point represents independent transfections. At least 100 cells were counted over three separate trials; data points represent individual transfections and bars show means ± SEM, n= 3-4. An ordinary one-way ANOVA with Dunnett’s multiple comparisons test was used to compare the percentage of cells with foci in each condition to cells expressing FL Kapβ2. **, p<0.005; ***, p<0.0005; ****, p<0.0001. (**C**) Quantifying the ratio of GFP signal coming from the cytoplasm and the nucleus shows that H1-H13, H8-E, and H9-E promote nuclear FUS localization to a level that is similar to that of FL Kapβ2. Η1-Η10 and H12-E however, are not as effective. Each point represents a single cell, and bars represent means ± SEM, n > 150 for each condition. An ordinary one-way ANOVA with Dunnett’s multiple comparisons test was used to compare the nuclear GFP-FUS signal in each condition to cells expressing FL Kapβ2. *, p<0.05; **, p<0.005; ***, p<0.0002; ****, p<0.0001.

We next assessed the ratio GFP-FUS signal coming from the cytoplasm and the nucleus to measure how proficient each construct is as a NIR (**Figure 7C**). As expected, expression of Kapβ2 promoted nuclear localization of GFP-FUS significantly more than expression of mCherry alone (**Figure 7C**). Conversely, Kapβ2^W460A:W730A^ did not increase the fraction of GFP-FUS in the nucleus (**Figure 7C**). Similar to their inability to reduce cytoplasmic FUS foci, H1-H10 and H12-E were also unable to restore nuclear localization of FUS as well as Kapβ2 (**Figure 7C**). By contrast, H8-E promoted nuclear localization of GFP-FUS as effectively as Kapβ2.

Curiously, H1-H13 and H9-E also promoted nuclear localization of GFP-FUS as well as Kapβ2 (**Figure 7C**). This finding was unexpected, as in yeast these truncations could only reduce cytoplasmic foci (**Figure 3B,C**). However, one important difference between the yeast and HeLa cell models is that yeast lack Kapβ2 and FUS. Indeed, yeast utilize Kap104 to perform Kapβ2-like transport activity (12,13,105). Although Kap104 can weakly bind to the PY-NLS of FUS, Kap104 preferentially binds basic PY-NLS sequences, whereas FUS harbors a hydrophobic PY-NLS (13, 14). Indeed, any *in vitro* binding capacity Kap104 may retain for FUS is insufficient for chaperone activity in yeast when FUS is overexpressed (**Figure 2**) (16, 90). By contrast, HeLa cells harbor endogenous Kapβ2. Thus, even low levels of truncated Kapβ2 variants may be able to collaborate with endogenous Kapβ2 to effectively reduce formation of cytoplasmic FUS foci and promote nuclear localization of FUS (**Figure 7**). Indeed, in HeLa cells, there is a noticeable increase in the cytoplasmic mCherry signal for H1-H10, H1-H13, H9-E, and H12-E, suggesting these proteins may be defective for nuclear import (**Figure 7A**). The increased levels of cytoplasmic truncated Kapβ2 suggest that all truncations except H8-E are deficient in their ability to traverse in and out of the nucleus, but retain some inherent chaperone activity in the cytoplasm that cooperates with endogenous Kapβ2. In yeast, this cytoplasmic chaperone activity manifests as a diffuse cytoplasmic FUS signal as Kap104 does not import FUS into the nucleus (13,89,90). In HeLa cells, cytoplasmic chaperone activity may help liberate FUS from cytoplasmic foci, making it available to the endogenous transport-competent Kapβ2.

Our findings suggest that H8-E is a robust chaperone, performing as well as FL Kapβ2 in several contexts, including: (1) preventing FUS toxicity and promoting FUS nuclear localization in yeast; (2) preventing FUS fibril formation *in vitro*; (3) dispersing FUS liquid condensates in vitro; and (4) reducing cytoplasmic FUS foci and restoring nuclear localization of FUS in human cells. Our data demonstrate that the N-terminal H1-H7 region of Kapβ2 are not required for significant chaperone functionality. Nonetheless, the N-terminal H1-H7 region enables optimal disaggregation of solid phase FUS condensates, indicating a functional role in disaggregation. Our findings definitively map the molecular chaperone and nuclear-import activities of Kapβ2 to H8-H20.

## Discussion

Kapβ2 is a NIR that also demonstrates specific chaperone activity against its cargo *in vitro*, in yeast, in human cells, and in fly models (16). The structure and localization of Kapβ2 in cargo-bound and unbound states have been extensively characterized (7,8,12,79,81,84). However, it has remained unclear precisely how the structure of Kapβ2 is directly related to its chaperone activity. Here, we designed truncated variants of Kapβ2 that eliminate its binding to specific epitopes of the FUS PY-NLS to determine which regions of Kapβ2 are most critical for chaperoning cargo and importing cargo into the nucleus.

In a yeast model system where the Kapβ2 cargo protein FUS is toxic (89–91), FL Kapβ2 mitigates FUS toxicity. H8-E, the only Kapβ2 truncation that includes all three PY-NLS epitope-binding regions, can also suppress FUS toxicity in this model. When GFP-tagged FUS is expressed in yeast, it forms abundant cytoplasmic aggregates and is depleted from the nucleus. Expression of FL Kapβ2 resolves these cytoplasmic aggregates and localizes FUS to the nucleus. In line with the results from yeast toxicity assays, H8-E also eliminates cytoplasmic FUS foci and promotes nuclear FUS localization. Indeed, H8-E was the only Kapβ2 truncation variant tested here to phenocopy Kapβ2 in yeast. None of the other Kapβ2 truncation constructs could mitigate FUS aggregation *and* toxicity.

Interestingly, truncations H1-H13 and H9-E, which did not mitigate FUS toxicity in yeast, displayed an intermediate phenotype where cytoplasmic FUS foci were partially dissolved without robust nuclear import. Previous work has established that dissolving cytoplasmic FUS aggregates with potentiated variants of the AAA+ disaggregase Hsp104 without promoting nuclear import can suppress FUS toxicity (91,94,96,106). However, the partial decrease in cytoplasmic FUS foci conferred by H1-H13 or H9-E is insufficient to suppress FUS toxicity. Thus, our data suggest that truncated variants of Kapβ2 that contain only a portion of the known PY-NLS-contacting residues have a diminished capability to antagonize cytoplasmic FUS aggregation, restore nuclear localization, and prevent cargo-related toxicity in yeast.

We also assessed the ability of truncated Kapβ2 proteins to directly chaperone FUS at the pure protein level. Kapβ2 potently prevents and reverses FUS aggregation in this setting (16). H8-E, which counters FUS toxicity and aggregation in yeast as well as FL Kapβ2, also chaperones FUS *in vitro*, although its ability to disaggregate FUS fibrils is mildly impaired compared to that of FL Kapβ2. H1-H13, on the other hand, only chaperones FUS at very high concentrations, and was unable to completely prevent or reverse FUS aggregation. H8-H16 can weakly prevent FUS aggregation, but its chaperone activity is substantially lower than even that of H1-H13. Surprisingly, although H9-E makes partial contact with the PY-epitope of the PY-NLS of FUS, the protein was inactive *in vitro*. However, the yield from the purification of H8-H16 and H9-E was low compared to H1-H13, H8-E, or Kapβ2, indicating that H8-H16 and H9-E may be less stable, which could affect chaperone activity. Nevertheless, these data indicate that HEAT repeat 8 plays a critical role for *in vitro* chaperone activity, and that the entire FUS-PY-NLS-binding region of Kapβ2 enables chaperone activity. We also assessed the ability of Kapβ2 and truncated variants to dissolve FUS:RNA liquid condensates. Here too, H1-H13 has ∼50% FL Kapβ2 activity, whereas FL Kapβ2 and H8-E rapidly dissolve FUS:RNA liquid condensates, significantly reducing both their number and total area.

Why is H8-E, which performs as well as Kapβ2 in all other respects, mildly deficient (i.e. ∼2-fold increase in EC_50_) in disaggregating preformed FUS fibrils? This finding suggests that the N-terminal H1-H7 region makes a contribution toward optimal disaggregation of FUS fibrils. We suggest that H1-H7 may make secondary contacts with the FUS PrLD after Kapβ2 has engaged the FUS PY-NLS, which help break the intermolecular contacts that hold FUS fibrils together (6,16,19). These contacts may be akin to interactions that NIRs make with FG-rich nucleoporins (Nups) during passage across the nuclear-pore complex (107–110). Indeed, the N-terminal HEAT repeats of a related NIR, Kapβ1, make contacts with FG-rich Nups (107). Moreover, a related nuclear-export factor, CRM1, may prevent FG-rich Nup condensation in the cytoplasm (111). Regardless, H8-E can still disaggregate FUS fibrils, albeit less efficiently than FL Kapβ2, indicating that residues beyond H1-H7 can also break contacts between FUS PrLDs that preserve fibril integrity.

A critical element in the structure of Kapβ2 is an extended acidic loop in H8. In the cytoplasm, where Ran is primarily in its GDP-bound state, the H8 loop is disordered and the Kapβ2 cargo-binding surface is available to interact with cargo (1,7,80,84). When Kapβ2 is bound to Ran-GTP in the nucleus, the H8 loop repositions to occupy the internal cargo-binding surface of Kapβ2, displacing the loaded cargo (112). Thus, this extended loop is necessary for cargo unloading in the presence of Ran-GTP (81, 112). The extended loop is not, however, required for Kapβ2 disaggregation activity (16). Indeed, Kapβ2 with a truncated acidic loop (Kapβ2^TL^, in which residues 341-362 are replaced by a 7-residue GGSGGSG linker) still acts as a chaperone *in vitro* to prevent and reverse FUS aggregation, and Kapβ2^TL^ prevents and reverses FUS recruitment to SGs in cells (16, 112). Thus, Kapβ2 chaperone activity can be separated from its role in nuclear import. Nonetheless, Kapβ2^TL^ would have limited ability to restore FUS to the nucleus and mitigate any loss of FUS nuclear function.

We also assessed the activity of Kapβ2 and truncated variants in human cells. Overexpression of GFP-FUS in HeLa cells results in formation of cytoplasmic FUS foci. However, when mCherry-FL-Kapβ2 or -H8-E is co-expressed, GFP-FUS is transported into the nucleus and the proportion of cells with cytoplasmic GFP-FUS foci decreases. Coexpression of H1-H13 or H9-E also reduces the number of cells with cytoplasmic FUS foci, despite displaying reduced chaperone activity *in vitro*. Surprisingly, despite being unable to promote nuclear localization in yeast, H1-H13 and H9-E do promote nuclear localization of FUS in human cells. We therefore suggest that these Kapβ2 truncations may help endogenous Kapβ2 to antagonize formation of cytoplasmic FUS foci, allowing endogenous Kapβ2 to promote FUS nuclear localization.

Our findings establish that the entire FUS-binding region of Kapβ2 is required for effective chaperone and nuclear-import activity. Cargo recognition requires the eighth HEAT repeat, but the residues contained in this region are not sufficient for functionality. The importance of H8 here is consistent with previous work, as H8 contains the residues that bind to the PY epitope of the PY-NLS, and this epitope contributes to the tight binding between Kapβ2 and cargo (1, 14). However, our work indicates that binding to non-PY-epitopes is also important and supports overall Kapβ2 function. Indeed, Kapβ2 relies on a large number of low-affinity interactions to chaperone cargo (12,19,23,113). Moreover, these data agree with genetic studies that have identified mutations in all three PY-NLS epitopes of FUS as causing ALS (14,77,114,115).

Similar mutations in the PY-NLS of hnRNPA1 and hnRNPA2 also cause degenerative disease (40,41,116,117). We suggest that Kapβ2 chaperone activity is achieved by making an extensive network of both strong and weak interactions with cargo, displacing cargo:cargo interactions in favor of Kapβ2:cargo interactions. Mutating just two residues in Kapβ2 that engage cargo as in the Kapβ2^W460A:W730A^ variant inactivates Kapβ2. Interestingly, H8-H16 and H9-E, each of which contains both Trp residues that are mutated to Ala in the Kapβ2^W460A:W730A^ mutant, are inactive in yeast despite being well expressed. Thus, the specific interactions mediated by W460 and W730 are important but not sufficient for chaperone activity.

Can truncated Kapβ2 proteins be used to supplement endogenous levels of Kapβ2 function? In models of disease where Kapβ2 cargo proteins aggregate, increasing expression of Kapβ2 can restore cargo-protein functionality (16). Thus, boosting Kapβ2 activity is an intriguing therapeutic approach (6,25–28,118). Kapβ2 is a large protein, and thus delivering smaller portions of the protein may be a more tractable approach. Recent advances in AAV-mediated delivery of genetic material for exogenous protein production have made the prospect of gene therapy more realistic (119–121). However, AAVs are limited by their DNA-packaging capacity, which restricts the size of the protein they can express. Of the two AAV-based treatments approved by the United States Food and Drug Administration (FDA), the molecular weight of the largest protein to be expressed is ∼65 kDa (122, 123). Kapβ2 is ∼100 kDa, whereas H8-E is ∼70 kDa.

Thus, it may be that with further technological improvements to AAV-based approaches, H8-E or even FL Kapβ2 could be delivered exogenously to treat diseases where nucleocytoplasmic trafficking of PY-NLS cargo is impaired, including ALS/FTD, MSP, oculopharyngeal muscular dystrophy, hereditary motor neuropathy, and related disorders (39-41,124-127).

## Experimental Procedures

**Table 1.** Key Resources.

| REAGENT or RESOURCE | SOURCE | IDENTIFIER |
| --- | --- | --- |
| **Antibodies** |  |  |
| Rabbit polyclonal anti-FUS | Bethyl | Cat# A300-302A; RRID: AB_309445 |
| Mouse monoclonal anti-Flag (M2) | Sigma-Aldrich | Cat# F3165; RRID: AB_259529 |
| Anti-Kapβ2 monoclonal antibody | Novus | Cat# NB600-1397 |
| Rabbit monoclonal anti-alpha Tubulin antibody | Abcam | Cat# ab184970 |
| Rat monoclonal anti-Tubulin antibody | Abcam | Cat# ab6160 |
| Anti-GFP polyclonal antibody | Sigma-Aldrich | Cat# G1544 |
| Anti-mCherry polyclonal antibody | Abcam | Cat# ab167453 |
| Goat anti-rabbit secondary antibody | Li-cor | Cat# 926-68071; RRID: AB_10956166 |
| Goat anti-mouse secondary antibody | Li-cor | Cat# 926-32210; RRID: AB_621842 |
| Goat anti-rat secondary antibody | Li-cor | Cat# 926-32219; RRID: AB_1850025 |
| **Bacterial and Virus Strains** |  |  |
| *E. coli* BL21-CodonPlus(DE3)-RIL | Agilent | Cat# 230245 |
| *E. coli* XL 10-Gold cells | Agilent | Cat# 200314 |
| One Shot TOP10 Chemically Competent *E. coli* | Invitrogen | Cat# C404010 |
| **Chemicals, Peptides, and Recombinant Proteins** |  |  |
| PfuUltra II Fusion HS DNA Polymerase | Agilent | Cat# 600672 |
| DpnI | NEB | Cat# R0176S |
| Gateway BP Clonase II Enzyme Mix | Invitrogen | Cat# 11789013 |
| Gateway BP Clonase II Enzyme Mix | Invitrogen | Cat# 11791019 |
| NEBuilder<sup>®</sup> HiFi DNA Assembly Master Mix | NEB | Cat# E2621S |
| Ni-NTA agarose | QIAGEN | Cat# 30230 |
| Glutathione Sepharose® 4 Fast Flow | GE Healthcare | Cat# GE17-5132-01 |
| cOmplete™, EDTA-free Protease Inhibitor Cocktail | Roche | Cat# 11873580001 |
| Hoechst 33342 stain | Invitrogen | Cat# H3570 |
| Lipofectamine™ 2000 Transfection Reagent | Invitrogen | Cat# 11668027 |
| VECTASHIELD® Antifade Mounting Medium with DAPI | Vector Laboratories | Cat# H-1200-10 |
| DMEM, high glucose, pyruvate | Gibco | Cat# 11995065 |
| Fetal Bovine Serum | Cytiva | Cat# SH30910.03 |
| Penicillin-Streptomycin | Gibco | Cat# 15140122 |
| HisTrap™ Fast Flow Crude | Cytiva | Cat# 17528601 |
| Novex AcTEV Protease | Fisher Scientific | Cat# 12-575-015 |
| Cy3 NHS Ester | Cytiva | Cat# PA13101 |
| **Critical Commercial Assays** |  |  |
| QuikChange Site-Directed Mutagenesis Kit | Agilent | Cat# 210518 |
| **Experimental Models: Organisms/Strains** |  |  |
| *S. cerevisiae*: W303aΔhsp104 (MATa, can1-100, his3-11, 15, leu2-3, 112, trp1-1, ura3-1, ade2-1, hsp104::KanMX) | Ref. (128) | A3224 |
| *S. cerevisiae*: W303aΔhsp104-pAG303GAL-FUS | Ref. (91) | N/A |
| *S. cerevisiae*: W303aΔhsp104-pAG303GAL-FUS-GFP | Ref. (91) | N/A |
| **Experimental Models: Cell Lines** |  |  |
| Human: HeLa | ATCC | Cat# CCL-2;<br>RRID:CVCL_0030 |
| **Recombinant DNA/RNA** |  |  |
| GST-TEV-FUS<sup>WT</sup> | Ref. (90) | N/A |
| GST-TEV-Kapβ2 | Ref. (16) | N/A |
| His-SUMO-Kapβ2 (full length and truncations) | This paper | N/A |
| pAG416Gal-Kapβ2 (FL and truncations) | This paper | N/A |
| mCherry2-C1 | N/A | Addgene Cat# 54563 |
| mCherry2-N1 | N/A | Addgene Cat# 54517 |
| pEGFP-C1-FLAG | Ref. (129) | GenBank U55763.1 |
| mCherry2-Kapβ2 (FL and truncations) | This paper | N/A |
| pEGFP-FUS<sup>WT</sup> | This paper | N/A |
| pTHMT/FUS<sub>WT</sub> | Ref. (20) | N/A |
| U<sub>40</sub> RNA: 5'- rUrUrU rUrUrU rUrUrU<br>rUrUrU rUrUrU rUrUrU rUrUrU rUrUrU<br>rUrUrU rUrUrU rUrUrU rUrUrU rUrUrU<br>rU/3AmMO/ -3' | IDT | N/A |
| **Software and Algorithms** |  |  |
| ImageJ | Ref. (130) | https://imagej.nih.gov/ij/ |
| GraphPad Prism | GraphPad | Version 9.1.1 |
| MATLAB | Mathworks |  |
| NIS-Elements Ar Package | Nikon |  |

### Plasmids

Kapβ2 truncations were cloned in the following manner: H1-H4, H1-H10, and H1-H13 were created by inserting stop codons into a GST-TEV-Kapβ2 plasmid using the following primers with their respective reverse complements:

5’-GACAGTGATGTTTTAGATCGTTAGTAGAACATCATG-3’;

5’-CAGCCGCCAGACACGTAGTAGAAGCCATTAATG-3’; and

5’-GGATTCCTTCCGTGATGAGAACCTGTGTATCAG-3’. H8-E, H9-E and H12-E were cloned into a GST-TEV-Kapβ2 plasmid (1, 16) using the following primers with their respective reverse complements:

5’-CAGGGCGGATCCAAGGCTTTAGAAGCCTGTGAATTTTGG-3’;

5’-CAGGGCGGATCCAAGGCTGCTGCCCTGGATGTTCTT-3’; and

5’-CAGGGCGGATCCAAGTACCTTGCTTATATACTTGATACCCTG-3’. Truncated constructs were then shuttled into a Gateway entry vector, pDONR221-ccdB, using Gateway BP reactions (Invitrogen). H8-H16 was generated at this point using the following primer to introduce a stop codon at the end of H16:

3’-GGGGACCACTTTGTACAAGAAAGCTGGGTCCTAAGGCTGCATCTCTATACCCATTTG-5’.

The entry clones were then used to shuttle Kapβ2 truncations into pAG416GAL-ccdB via Gateway LR reactions for yeast expression. FLAG-tagged constructs were created by using primers to add the following sequence upstream of the stop codon for each open-reading frame (ORF): 5’-GACTACAAAGACGATGACGACAAG-3’.

To create plasmids for expression in mammalian cells, FUS and Kapβ2 were cloned into pEGFP-C1 and mCherry2-C1 (Addgene, #54563), respectively. The FUS ORFs were amplified with the following overhang sequences: 5’-GGACGAGCTGTACAAG-3’;

3’-GTAATCAGATCTGAGTCCGG-5’. Kapβ2 ORFs were amplified with the following overhang sequences: 5’-GGACGAGCTGTACAAGTCT-3’; 3’-CTCGAGATCTGAGTCCGG-5’. For both ORFs, the vector backbone was cut with BspEI and the linearized backbone and amplified inserts were purified using QIAquick Gel Extraction Kit (Qiagen) and assembled using Gibson cloning (NEB).

To create plasmids for recombinant expression of Kapβ2 plasmids, Kapβ2 ORFs were cloned into a pE-SUMOpro Amp vector (LifeSensors) using the following strategy: first, the empty pE-SUMOpro Amp vector was cut using BsaI enzyme (NEB). ORFs were amplified with primers that added 5’-CGCGAACAGATTGGAGGT-3’ upstream of the start of the ORF, and added 5’-CTAGAGGATCCGAATTC-3’ downstream of the end of the ORF. The cut vector and amplified ORFs were purified using a QIAquick Gel Extraction Kit (Qiagen), and assembled using Gibson cloning (NEB).

All constructs were confirmed by DNA sequencing.

### Yeast Strains and Media

Yeast were the isogenic strain W303aΔ*hsp104* (91). The yeast strain W303aΔhsp104-pAG303GAL-FUS^WT^ and W303aΔhsp104-pAG303GAL-FUS^WT^-GFP have been described previously (91, 94). Media was supplemented with 2% glucose, raffinose, or galactose as specified.

### Yeast Transformation and Spotting Assays

Yeast transformations were performed using standard polyethylene glycol and lithium acetate procedures (131). For the spotting assays, yeast were grown to saturation in raffinose-supplemented dropout media overnight at 30 °C. The saturated overnight cultures were normalized to an OD_600_ of 2.0 (A_600nm_ = 2.0) and serially diluted three-fold. A 96-bolt replicator tool (frogger) was used to spot the strains in duplicate onto both glucose and galactose dropout plates. These plates were grown at 30 °C and imaged after 72 h to assess suppression of FUS toxicity. Each spotting assay shown is representative of at least three biological replicates.

### Western Blots

For yeast cells, transformed Kapβ2 variants and controls were grown overnight in raffinose media. The overnight cultures were diluted to an OD_600_ of 0.3 (A_600nm_ = 0.3) and grown in galactose-supplemented media at 30 °C for 5 h. Samples were then normalized to an OD_600_ of 0.6 (A_600nm_ = 0.6) and centrifuged at 4000 x g for 5 min. The pelleted cells were resuspended in 0.1 M NaOH for 5 min and then pelleted again and resuspended in 1x SDS sample buffer. For HeLa cells, ∼2-2.5 x 10^5^ cells were seeded and transfected with GFP-tagged FUS and either mCherry-tagged Kapβ2 variants or an empty vector expressing mCherry. After 24 h, cells were washed once with PBS, then resuspended in RIPA lysis buffer (150 mM NaCl, 1% Triton X-100, 1% sodium deoxycholate, 0.1% SDS, 25 mM Tris-HCl pH 7.6) supplemented with protease inhibitors and 1 mM PMSF. Cells were then sonicated and centrifuged at 4 °C for 10 minutes at 10,000 x g, and the cell lysate was mixed with 1x SDS sample buffer.

The samples were then boiled and separated by SDS-PAGE (4–20% gradient, Bio-Rad) and transferred to a PVDF membrane. The following primary antibodies were used: anti-FUS polyclonal (Bethyl Laboratories), anti-Kapβ2 monoclonal (Novus), anti-alpha Tubulin monoclonal (Abcam: ab184970 for yeast; ab6160 for human cells), anti-FLAG monoclonal (Sigma-Aldrich), anti-GFP polyclonal (Sigma-Aldrich), anti-mCherry polyclonal (Abcam). Three fluorescently labeled secondary antibodies were used: anti-rabbit (Li-Cor), anti-rat (Li-Cor), and anti-mouse (Li-Cor). Blots were imaged using a LI-COR Odyssey FC Imaging system.

### HeLa Cell Culture and Transfections

HeLa cells were maintained in Dulbecco’s modified Eagle’s medium, high glucose (Gibco) containing 10% fetal bovine serum (HyClone) and 1% penicillin-streptomycin (Gibco). Cells were plated in 6-well plates at a density of 2-2.5×10^5^ cells/plate 24 h before transfection. Cells were transfected with 1 µg total DNA and 3 µL Lipofectamine 2000 (Invitrogen). Media was changed 4 h post-transfection to standard maintenance media and cells were harvested for microscopy or western blot 24 h post-transfection.

### Fluorescence Microscopy

For yeast microscopy, W303aΔhsp104-pAG303GAL-FUS^WT^-GFP yeast were transformed with the indicated Kapβ2 variant or vector control. Microscopy samples were grown and induced as they were for immunoblotting. Prior to imaging, cells were treated with Hoechst 33342 stain (0.1 mg/mL). All cells were imaged at 100x magnification using a Leica-DMIRBE microscope or the EVOS M5000 Imaging System. Analysis of cells was performed in ImageJ (132). For each sample, at least 100 cells were quantified in at least three independent trials.

For HeLa cell microscopy, transfected HeLa cells were fixed with 2% formaldehyde for 30 min at RT, followed by treatment with Triton X-100 for 6 min to permeabilize cells. Coverslips were then assembled using VECTASHIELD^®^ Antifade Mounting Medium with DAPI (Vector Laboratories) and sealed before imaging. Images were taken at 100x magnification using the EVOS M5000 Imaging System (ThermoFisher) and processed using ImageJ (132). At least 100 cells were counted for each condition across three independent trials.

### Protein Purification

All proteins were expressed and purified from *E. coli* BL21-CodonPlus(DE3)-RIL cells (Agilent) and purified under native conditions. GST-TEV-FUS^WT^ was purified as described (Sun et al., 2011). Briefly, *E. coli* cells were lysed by sonication on ice in PBS pH 7.4 with protease inhibitors (cOmplete, EDTA-free, Roche Applied Science). The protein was purified over Glutathione Sepharose 4 Fast Flow (GE Healthcare) and eluted from the beads using 50 mM Tris-HCl pH 8, 20 mM trehalose, and 20 mM glutathione. 6xHis-MBP-FUS was purified as described (20). Briefly, *E. coli* cells were lysed by sonication in lysis buffer (1 M KCl, 1 M Urea, 50 mM Tris-HCl pH 7.4, 10 mM imidazole, 1.5 mM β-mercaptoethanol, 1% NP-40, 5% glycerol, supplemented with protease inhibitors). The protein was purified over a HisTrap HP column (GE), using an AKTA pure 25M FPLC system (GE), and was eluted using an imidazole gradient into elution buffer (1M KCl, 1 M Urea, 50 mM Tris-HCl pH 7.4, 1.5 mM β-mercaptoethanol, 5% glycerol, 500 mM imidazole), and the appropriate fractions were pooled and stored in 30% glycerol at 4 °C.

To purify full-length and truncated Kapβ2 proteins, *E. coli* cells transformed with the appropriate plasmid were grown at 37 °C in LB supplemented with 2% glucose. Once cells reached an OD_600_ of ∼0.6, expression was induced overnight at 15 °C with 0.2-1.0 mM isopropyl β-d-1-thiogalactopyranoside (IPTG). Cells were pelleted and resuspended in binding buffer (50 mM Tris-HCl, pH 7.5, 100 mM NaCl, 20% glycerol, 10 mM imidazole, 2.5 mM β-mercaptoethanol, supplemented with protease inhibitors), then lysed by sonication. Cell lysate was separated by spinning at 16,000 rpm at 4 °C for 20 min. Cell lysate was then loaded onto Ni-NTA resin (Qiagen) and washed with binding buffer. Protein was eluted with elution buffer (50 mM Tris-HCl, pH 7.5, 100 mM NaCl, 20% glycerol, 200 mM imidazole, 2.5 mM β-mercaptoethanol) and buffer exchanged into a low imidazole buffer (20 mM imidazole pH 6.5, 75 mM NaCl, 2.5 mM β-mercaptoethanol, 20% glycerol). Protein was then cleaved at 30 °C overnight with Ulp1 at a 1:250 molar ratio in an Eppendorf ThermoMixer shaking at 300 rpm. Cleaved protein product was then bound to fresh Ni-NTA resin, and flow-through was collected and further purified on a HiTrap Q HP column (GE) using buffer A (20 mM imidazole pH 6.5, 75 mM NaCl, 1 mM EDTA, 2 mM DTT, 20% glycerol) and buffer B (20 mM imidazole pH 6.5, 1 M NaCl, 1 mM EDTA, 2 mM DTT, 20% glycerol). Protein fractions were then concentrated, snap-frozen in liquid nitrogen, and stored at -80 °C.

### FUS Aggregation Assays

Spontaneous FUS fibrillization (i.e., in the absence of preformed fibril seeds) was initiated as described previously (16). Briefly, 1 µg TEV protease was added to GST-TEV-FUS (3 μM) in FUS assembly buffer (50 mM Tris-HCl, pH 8, 200 mM trehalose, 1 mM DTT, and 20mM glutathione) (37,38,90). Spontaneous FUS fibrillization reactions were incubated at 25 °C with or without the indicated concentration of each Kapβ2 variant, and turbidity was used to assess fibrillization by measuring absorbance at 395nm. The absorbance was normalized to that of FUS plus buffer control to determine the relative extent of aggregation. Disaggregation assays were performed similarly, where FUS aggregation was initiated with TEV in the absence of Kapβ2. After 60 minutes of aggregation, equal volumes were added to each condition containing the appropriate ratios of buffer and Kapβ2 to reach the indicated final concentration.

To analyze data, the reading from the first timepoint was subtracted from every subsequent reading. Then, for each aggregation experiment, the maximum absorbance value for FUS alone was set as 100, and all other values were normalized to that standard by dividing the raw, background-subtracted value by the maximum value for FUS alone and multiplying by 100. For disaggregation experiments, the background-subtracted reading from the time point immediately before Kapβ2 addition was set to 100 for each condition, and subsequent readings were normalized to their own initial standard. For aggregation assays, the area under the curve (AUC) for the first 90 minutes of aggregation was used to calculate the IC_50_ by dividing the AUC for each condition by the AUC for FUS alone and multiplying by 100 to calculate a normalized percent aggregation. The same data treatment was used for disaggregation assays, where the AUC for the first 40 minutes after addition of Kapβ2 or buffer was used to calculate the EC_50_. AUC, IC_50_, and EC_50_ were calculated using GraphPad Prism by fitting data to a non-linear regression using a normalized response and a variable slope.

### Droplet Assays

U_40_ RNA was labeled as described previously (20). Briefly, unlabeled amine-modified RNA was combined with 0.1 mg Cy3-NHS in 100 mM sodium bicarbonate for overnight labeling. The RNA was precipitated with ethanol twice and stored at -20 °C as 100 μM long-term and 1 μM working stocks.

Droplet reactions were performed essentially as described previously with a few modifications (20). Purified 6xHis-MBP-FUS was buffer-exchanged by concentrating FUS in Amicon filters (Millipore Sigma) and diluting with 20 mM Na2HPO4 pH 8.0. FUS (1 μM) was combined with unlabeled U40 RNA (1 μM) and Cy3-U40 (10 nM) in 1X Cleavage Buffer (100 mM NaCl, 50 mM Tris pH 7.4., 1 mM DTT, and 1 mM EDTA pH 8.0.). The phase separation reaction was initiated by adding 5 U Novex AcTEV protease, which cleaved the MBP and His tags from the FUS protein. Reactions (200 μL) were incubated for 4 h in 8-well Nunc Lab-Tek chambers (Thermo Scientific #155361). Brightfield and Cy3 fluorescence images were captured on a Nikon Ti Eclipse wide-field microscope. Purified Kapβ2 protein was prepared as a 10X concentrate (2.5-10 μM) in 1X Cleavage Buffer and diluted to 1X Kapβ2 (250-1000 nM) in the phase separation reactions immediately after the 4-h timepoint.

Droplet area and number were quantified with a MATLAB script that used intensity thresholding to identify droplet ROIs (21). We defined the area density as the sum of the area of all droplet ROIs divided by the total imaging area, which estimates the fraction of surface area covered by droplets. Therefore, the area density is unitless. The number of droplet ROIs and the area density of each image were determined for each replicate, which were then used to calculate the droplet statistics before and after Kapβ2 addition.

## Data availability

All data are contained within the manuscript.

## Supporting information

This article contains supporting information (supplementary figures 1-4).

## Acknowledgments

We thank Katie Copley, Miriam Linsenmeier, JiaBei Lin, Zarin Tabassum, and Hana Odeh for feedback on the manuscript.

## Funding and additional information

C.M.F. is supported by NIH grants T32GM008275 and F31NS111870. K.R. is supported by NIH grants F31NS113439 and T32GM007231. S.M. is funded by the NIH grants RF1AG071326 and RF1NS113636 and the NSF grant PHY-1430124 via the Center of Physics of Living Cells. J.S. is supported by grants from Target ALS, the ALS Association, the Office of the Assistant Secretary of Defense for Health Affairs through the Amyotrophic Lateral Sclerosis Research Program (W81XWH-20-1-0242), the G. Harold & Leila Y. Mathers Foundation, Sanofi, and NIH grants R01GM099836, R21AG061784, and R21AG065854.

## Author Contributions

C.M.F., K.R., A.L., S.M. and J.S. designed experiments. C.M.F. and A.L. performed cloning and yeast experiments, and C.M.F. performed human cell and *in vitro* purification and aggregation experiments. K.R. performed *in vitro* MBP-FUS purification and FUS:RNA droplet experiments. C.M.F. and J.S. wrote the paper.

---

**Figure S1.**
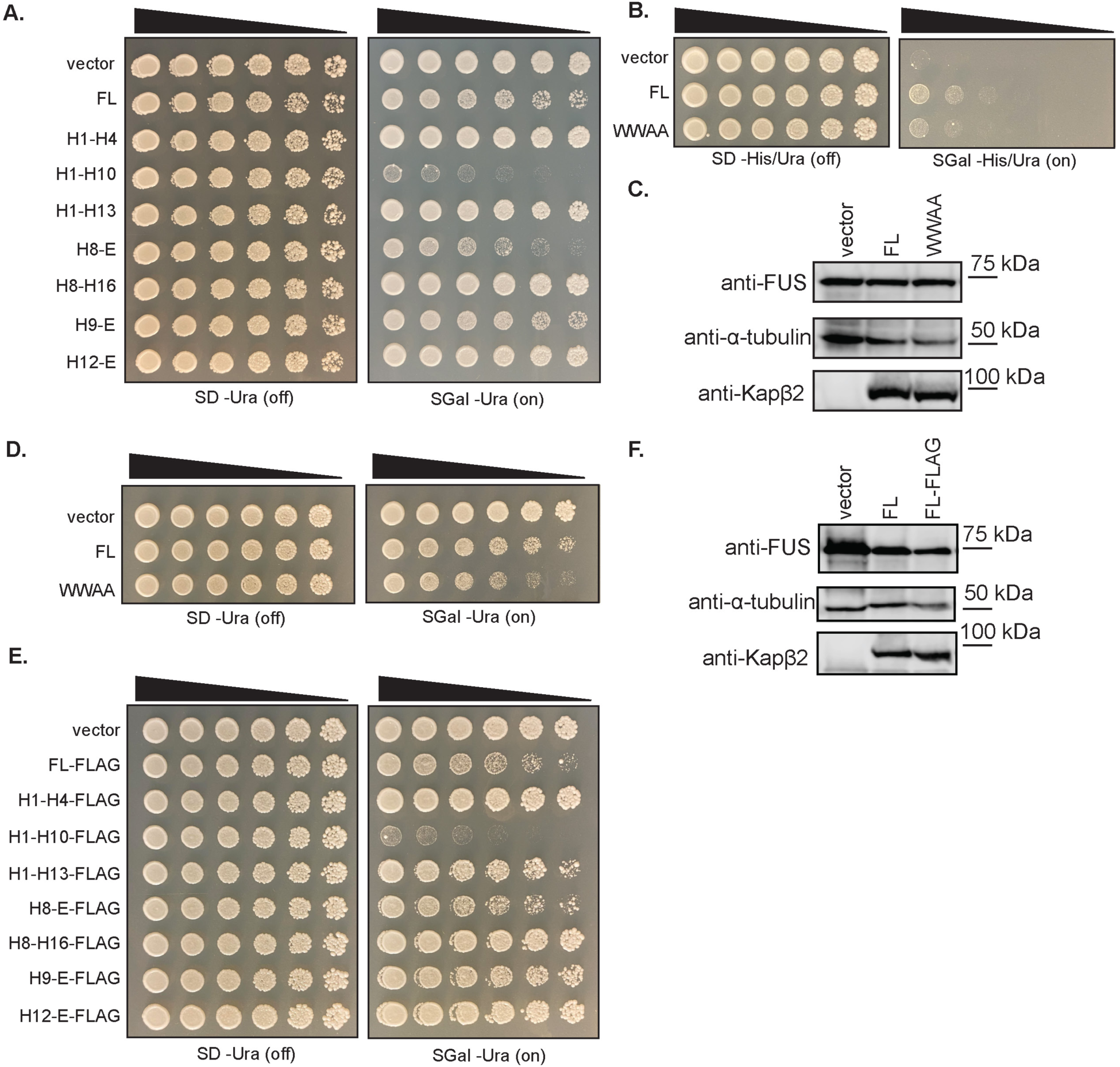
Kapβ2 and truncation variants are typically not toxic and robustly expressed in yeast. (**A**) Δ*hsp104* yeast were transformed with the indicated galactose-inducible Kapβ2 construct and serially diluted 3-fold onto agar plates with either glucose (expression off) or galactose (expression on) to assess toxicity of each Kapβ2 construct on its own. H1-H10 exhibits toxicity relative to vector, and FL Kapβ2, H8-E, and H9-E show slight toxicity. (**B**) Δ*hsp104* yeast with galactose-inducible FUS integrated into the strain were transformed with galactose-inducible FL Kapβ2 or Kapβ2^W460A:W730A^ (WWAA) and serially diluted 3-fold onto agar plates with either glucose (expression off) or galactose (expression on) as the carbon source. Media was prepared lacking histidine and uracil; the FUS plasmid contains genetic information for synthesizing histidine, and the Kapβ2 plasmid contains the genetic information for synthesizing uracil. In the absence of any Kapβ2, FUS expression is very toxic. This toxicity can be mitigated by the expression of full-length (FL) Kapβ2, but not the Kapβ2^W460A:W730A^ (WWAA) mutant. (**C**) A western blot probing yeast expressing FUS and either FL Kapβ2^WT^ or the WWAA mutant with an anti-Kapβ2 antibody shows that neither FUS nor Kapβ2 expression changes as a function of Kapβ2 variant expression relative to the α-tubulin loading control. (**D**) Δ*hsp104* yeast with no disease-associated protein integrated into the strain were transformed with either vector, FL Kapβ2 or the Kapβ2^W460A:W730A^ (WWAA) mutant and serially diluted 3-fold onto agar plates with either glucose (expression off) or galactose (expression on) as the carbon source to test for toxicity of WWAA on its own. WWAA is slightly more toxic than the WT FL protein. (**E**) Δ*hsp104* yeast with no disease-associated protein integrated into the strain were transformed with the indicated galactose-inducible FLAG-tagged Kapβ2 truncation and serially diluted 3-fold onto agar plates with either glucose (expression off) or galactose (expression on) as the carbon source to test for toxicity of each truncation on its own. As with the untagged variants, H1-H10-FLAG is more toxic relative to vector. H8-E-FLAG and H9-E-FLAG show less intrinsic toxicity than their untagged counterparts. (**F**) Probing yeast expressing FUS and either tagged or untagged FL Kapβ2 with an anti-Kapβ2 antibody shows that, compared to the untagged protein levels, the addition of the FLAG-tag does not affect FL Kapβ2 expression.

**Figure S2.**
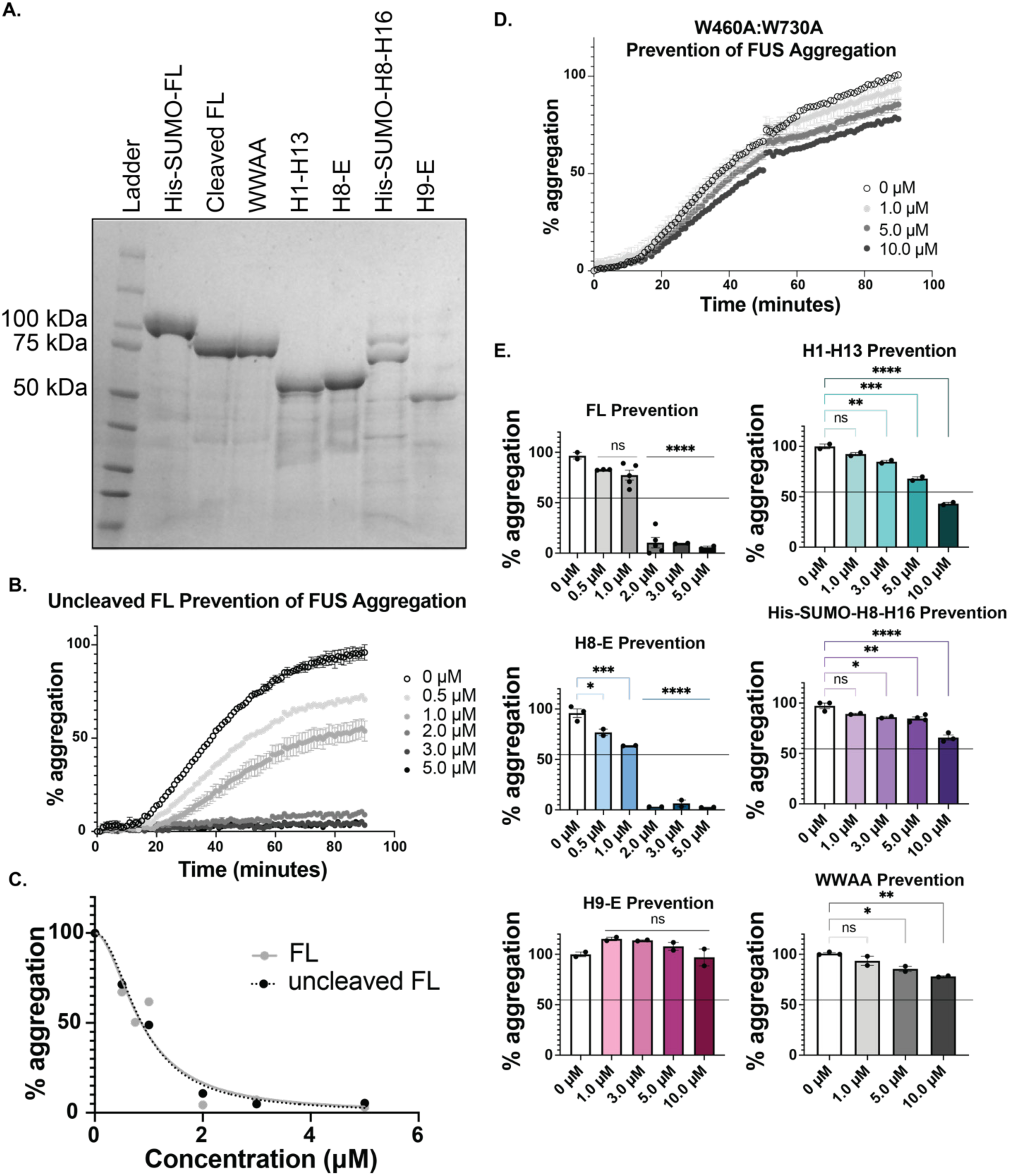
Kapβ2 and H8-E robustly suppress FUS aggregation. (**A**) 5 µg of purified His-SUMO-FL Kapβ2, cleaved FL Kapβ2, H1-H13, H8-E, His-SUMO-H8-H16, and H9-E were processed for SDS-PAGE and stained with Coomassie. (**B**) To measure the ability of His-SUMO-tagged Kapβ2 to prevent FUS aggregation, we incubated 3 μM FUS with 1 μg TEV protease and the indicated concentration of His-SUMO-tagged Kapβ2 and monitored turbidity at 395 nm. Increased turbidity indicates FUS self-assembly and each curve represents the mean of 2 trials ± SEM. For each experiment, the maximal value of FUS aggregation in the absence of Kapβ2 was set to 100 and all data for that trial were normalized to that value. (**C**) To compare uncleaved and cleaved FL Kapβ2 chaperone activity, we calculated the area under the curve (AUC) for each concentration and set the AUC for FUS alone as 100 to normalize all other conditions to produce percent aggregation. Overlaying the plot of concentration vs. aggregation (%) for cleaved and uncleaved FL Kapβ2 shows that the uncleaved protein works as well as cleaved Kapβ2 in preventing the aggregation of FUS, indicating that the His-SUMO tag itself neither hinders nor enhances Kapβ2 chaperone activity in this assay. (**D**) We assessed the ability of Kapβ2^W460A:W730A^ (WWAA) to chaperone FUS. We incubated 3 μM FUS with 1 μg TEV protease and the indicated concentration of Kapβ2^W460A:W730A^ and monitored turbidity at 395 nm. Each curve represents the mean of 2-3 trials ± SEM. The maximal value of FUS aggregation in the absence of Kapβ2 was set to 100 and all data for that trial were normalized to that value. (**E**) In Figure 4, we assessed the ability of FL and truncated Kapβ2 constructs to chaperone FUS. We incubated 3 μM FUS with 1 μg TEV protease and the indicated concentration of each Kapβ2 construct and monitored turbidity at 395 nm for 90 minutes. Here, only the final normalized absorbance reading (where the maximum value for FUS alone is set as 100) is plotted for each Kapβ2 construct at each of the indicated concentrations. Data points represent individual trials and bars represent means ± SEM. An ordinary one-way ANOVA with Dunnett’s multiple comparisons test was used to compare the final time point of each condition to FUS alone for that construct. *, p<0.03; **, p<0.008; ***, p<0.0003; ****, p<0.0001.

**Figure S3.**
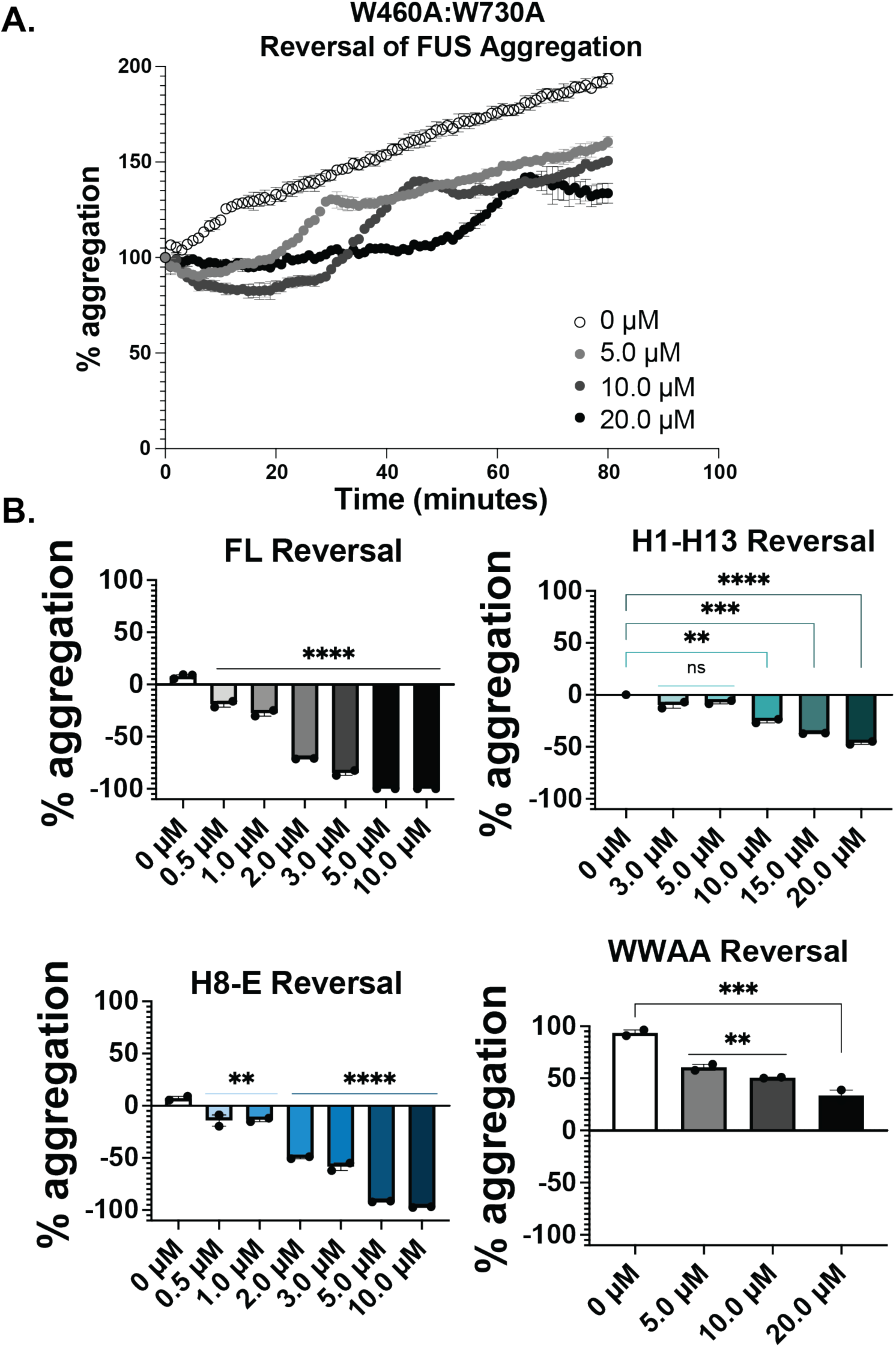
Kapβ2 and H8-E robustly reverse FUS aggregation. To measure the ability of Kapβ2^W460A:W730A^ (WWAA) to reverse FUS aggregation, we incubated 3 μM FUS with 1 μg TEV protease and monitored turbidity at 395 nm. After one hour of aggregation, the indicated amount of Kapβ2^W460A:W730A^ was added and turbidity was continually measured. Each curve represents the mean of 2-3 trials ± SEM. For each condition, the value of FUS aggregation immediately prior to adding Kapβ2 was set to 100 and all subsequent readings were normalized to that value. (**A**) Kinetics of FUS disaggregation by Kapβ2^W460A:W730A^ show that Kapβ2^W460A:W730A^ does not disaggregate FUS. (**B**) In Figure 5, we assessed the ability of FL and truncated Kapβ2 constructs to disaggregate FUS. We incubated 3 μM FUS with 1 μg TEV protease for 60 minutes before adding the indicated construct and measuring turbidity for 40 or 80 minutes. The difference between the starting value and the final reading for the normalized absorbance values (where timepoint 0 was set to 100) was measured for disaggregation reactions for each construct. Data points represent individual trials and bars represent means ± SEM. An ordinary one-way ANOVA with Dunnett’s multiple comparisons test was used to compare the final time point of each condition to FUS alone for that construct. **, p<0.005; ***, p<0.0006; ****, p<0.0001.

**Figure S4.**
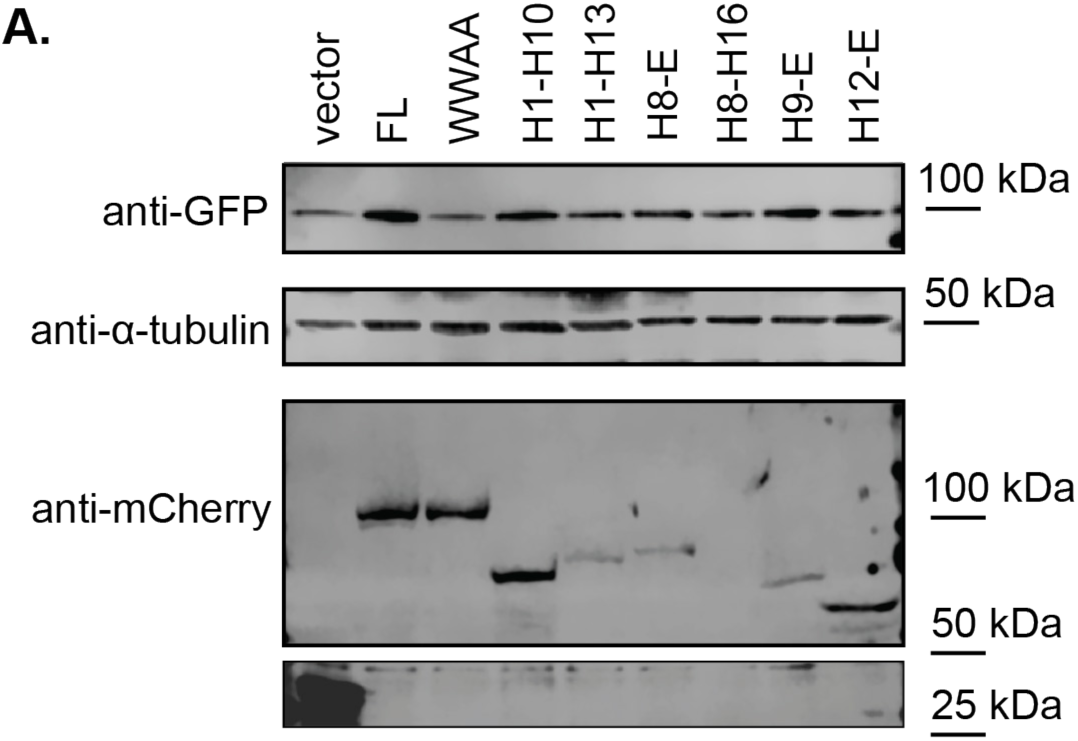
Kapβ2 and truncation variants are typically expressed in human cells. A western blot of lysates of HeLa cells transfected with GFP-FUS and either mCherry alone or mCherry-tagged FL Kapβ2 or the indicated construct. Probing with anti-GFP shows equal FUS-GFP expression. However, probing for anti-mCherry indicates that expression of select truncations is reduced (as for H1-H13, H8-E, and H9-E) or completely absent (H8-H16). α-tubulin is used as a loading control across samples.

